# Microbiome, resistome and mobilome of chlorine-free drinking water treatment systems

**DOI:** 10.1101/2022.12.08.519565

**Authors:** David Calderón-Franco, Francesc Corbera-Rubio, Marcos Cuesta-Sanz, Brent Pieterse, David de Ridder, Mark C. M. van Loosdrecht, Doris van Halem, Michele Laureni, David G. Weissbrodt

## Abstract

Drinking water treatment plants (DWTPs) are designed to remove physical, chemical, and biological contaminants. However, until recently, the role of DWTPs in minimizing the cycling of antibiotic resistance determinants has got limited attention. In particular, the risk of selecting antibiotic-resistant bacteria (ARB) is largely overlooked in chlorine-free DWTPs where biological processes are applied. Here, we combined high-throughput quantitative PCR and metagenomics to analyze the abundance and dynamics of microbial communities, antibiotic resistance genes (ARGs), and mobile genetic elements (MGEs) across the treatment trains of two chlorine-free DWTPs involving dune-based and reservoir-based systems. The microbial diversity of the water being treated increased after all biological unit operations, namely rapid and slow sand filtration (SSF), and granular activated carbon filtration. Both DWTPs reduced the concentration of ARGs and MGEs in the water by about 2.5 log gene copies mL^-1^, despite their relative increase in the disinfection sub-units (SSF in dune-based and UV treatment in reservoir-based DWTPs). The total microbial concentration was also reduced (2.5 log units), and none of the DWTPs were enriched for antibiotic resistant bacteria. Our findings highlight the effectiveness of chlorine-free DWTPs in supplying safe drinking water while reducing the concentration of antibiotic resistance determinants. To the best of our knowledge, this is the first study that monitors the presence and dynamics of antibiotic resistance determinants in chlorine-free DWTPs.

## 1. Introduction

Access to safe water and sanitation is a key Sustainable Development Goal (United Nations, 2015) and a central objective of the Water Action Decade (United Nations, 2018) of the United Nations. Drinking water treatment plants (DWTPs) are used to remove water contaminants and deliver safe water for consumption. The origin and nature of contaminants depend on several factors, such as the water source (Yu et al., 2018), geographical location (UNEP - United Nations Environment Programme, 2016), season (Kumpel et al., 2017), and type of anthropogenic activity in the water basin (Khatri and Tyagi, 2015). Contaminants can be divided into physical-chemical (*e.g*., suspended particles, iron, ammonia) and biological (*e.g*., pathogens, antimicrobial resistances – AMR) agents (United States Environmental Protection Agency, 2021).

The process configuration of DWTPs is mainly dictated by the water source, either groundwater or surface water. While groundwater is generally microbiologically safe, surface water may contain pathogenic organisms that must be eliminated (Smeets et al., 2009). Chemical disinfectants, such as chlorine, are usually applied to disinfect drinking water (i.e., inactivate pathogenic microorganisms) and/or to prevent microbial (re)growth in the distribution network (Sedlak and von Gunten, 2011). However, the use of disinfectants can generate by-products with mutagenic and carcinogenic effects (Rook, 1976) and selects for antibiotic-resistant bacteria (ARB) (Shi et al., 2013). A few countries (*e.g*., The Netherlands, Denmark, or Switzerland) ceased disinfectants use and rely on strict source-to-consumer production standards and engineering solutions for drinking water supply (Smeets et al., 2009). For the chlorine-free drinking water production from surface water, two main DWTP configurations (dune-based and reservoir-based; detailed description in Materials and Methods) are employed in the Netherlands. In both cases, a large fraction of the treatment consists of biological biofilm-based unit operations such as dune infiltration, rapid sand filtration (RSF), slow sand filtration (SSF), or granular activated carbon (GAC) filtration, which combine biological and physical-chemical processes. In these systems, chemical and biological water contaminants are converted by microbial communities (Mouchet, 1992; Tekerlekopoulou et al., 2013), which shape the microbiome of the drinking water that reaches consumers (Pinto et al., 2012). Therefore, the biological safety of the microbial communities harbored in DWTPs is of utmost importance for public health.

In contrast to wastewater environments (Calderón-Franco et al., 2022; Miłobedzka et al., 2022; Pallares-Vega et al., 2019), few studies focus on the fate and removal of ARGs and ARB in DWTPs. Biofilms are known reservoirs of ARB and antibiotic resistance genes (ARGs) (Balcázar et al., 2015). Little is known about the impact of biofilm-based DWTPs on the generation and/or persistence of ARB in drinking water. While antibiotic concentrations are very low or non-existent (Stackelberg et al., 2004), the generation of ARB in biofilms by horizontal gene transfer (HGT) is a well-known phenomenon (Farkas et al., 2013). Therefore, the effect of biofilms present in DWTPs operational-units on ARB development needs to be uncovered. To date, molecular studies of microbial communities, ARB, and ARGs in DWTPs have been limited by the low biomass concentration present in these systems for DNA extraction, sample collection logistics, and sampling standardization (Ma et al., 2017). The results often rely on either lab-scale experiments (Stange et al., 2019; Wan et al., 2019) or specific treatment processes such as biological activated carbon filters (Wan et al., 2021) or tertiary treatments such as chlorine, UV, or a combination of them (Destiani and Templeton, 2019; Shi et al., 2013). Therefore, information about how biological treatments affect the fate of ARGs and MGEs in full-scale DWTPs from an integral consideration of the treatment train and different geographical areas is missing.

Most integrative studies have been carried out in China (Hu et al., 2019; Jia et al., 2020, 2015; Su et al., 2018; Xu et al., 2016; Zhang et al., 2019, 2016), *i.e*., one of the largest antibiotic-producing and consuming countries world-wide (Huang et al., 2019). The studies have used qPCR (Hu et al., 2019; Su et al., 2018; Zhang et al., 2016), high-throughput qPCR (Xu et al., 2016) or sequencing methods like amplicon sequencing and metagenomics (Jia et al., 2020, 2015; Zhang et al., 2019) to investigate the concentration and richness of ARGs in full-scale DWTPs. Sevillano et al. (2020) have compared the effect of disinfection systems on antimicrobial resistance determinants on tap water samples in DWTPs from The Netherlands, UK, and USA. Yet, no detailed information about individual process units was provided. Moreover, none of these studies have combined qualitative (metagenomics) and quantitative (HT-qPCR) approaches for their analysis to get information about the total amount and diversity of AMR determinants in chlorine-free DWTPs.

In this work, we qualitatively and quantitatively resolve the role of chlorine-free DWTPs in the control of ARB and ARGs throughout the entire treatment train of two full-scale DWTPs in a low-antibiotic-consuming country. Specifically, we aim at deciphering how the biofilms in biological unit operations shape the resistome and mobilome of the drinking water. To do so, we compared the contribution of different methods for water storage, physical-chemical contaminant removal, and disinfection in one dune-based DWTP and one reservoir-based DWTP.

## 2. Material and methods

### 2.1. Sampling of two full-scale DWTPs

Water samples were collected from two different chlorine-free DWTPs supplying drinking water to the South Holland and Zeeland provinces in the Netherlands (Figure 1). The dune-based DWTP in this study infiltrates pre-treated river water into the sand dunes for storage and water quality improvement (e.g., disinfection). Subsequently, dune water is abstracted with wells and treated with pellet softening, powdered activated carbon (PAC), and rapid sand filtration (RSF) to remove hardness, organic (micro)contaminants, iron, and ammonium. Slow sand filtration (SSF) is deployed as a final disinfection step, as well as to ensure the biological stability of the water (i.e., remove trace nutrients). The reservoir-based DWTP in this study stores water in open reservoirs and uses a treatment train of coagulation-flocculation and RSF to eliminate physical-chemical contaminants, UV treatment for disinfection, and granular activated carbon (GAC) to remove organic contaminants (e.g., colour, odour, pesticides).

**Figure 1.**
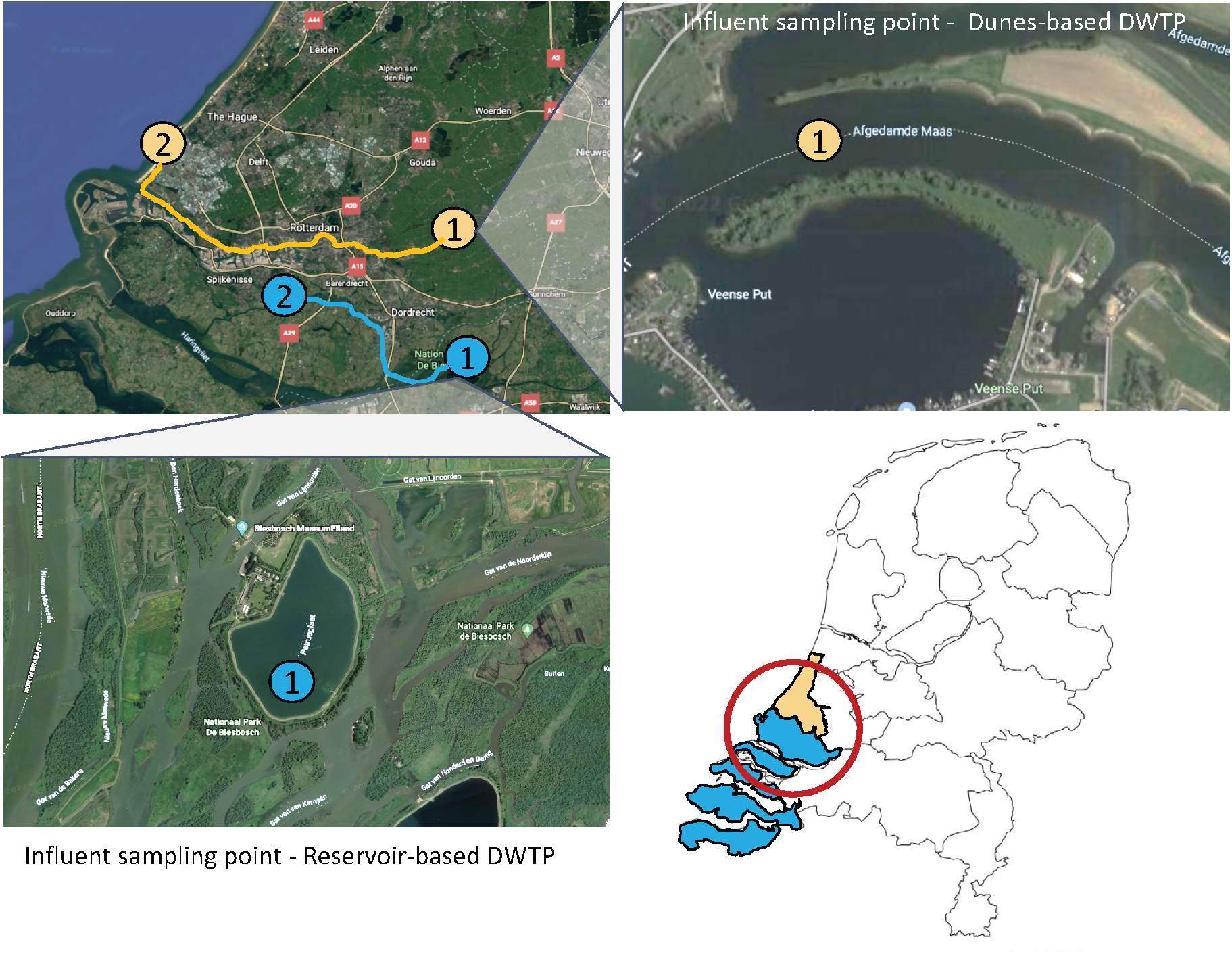
Geographic map of sampling sites. Numbering in the figure: (1) represents the river or reservoir from which the influent sample to the DWTP was taken, and (2) represents the location of the DWTP.

The first, dune-based DWTP(N 52° 7’ 1.9992; E 4° 18’ 23.9184) treats surface water from the Meuse River. The second, reservoir-based DWTP (N 51° 48’ 44.4132; E 4° 20’ 0.2112) processes surface water from the Meuse River as well (The Netherlands), after storage in a reservoir. Five sampling process stages per DWTP were targeted for water collection across their process stages (Figure 2, Table 1).

**Figure 2.**
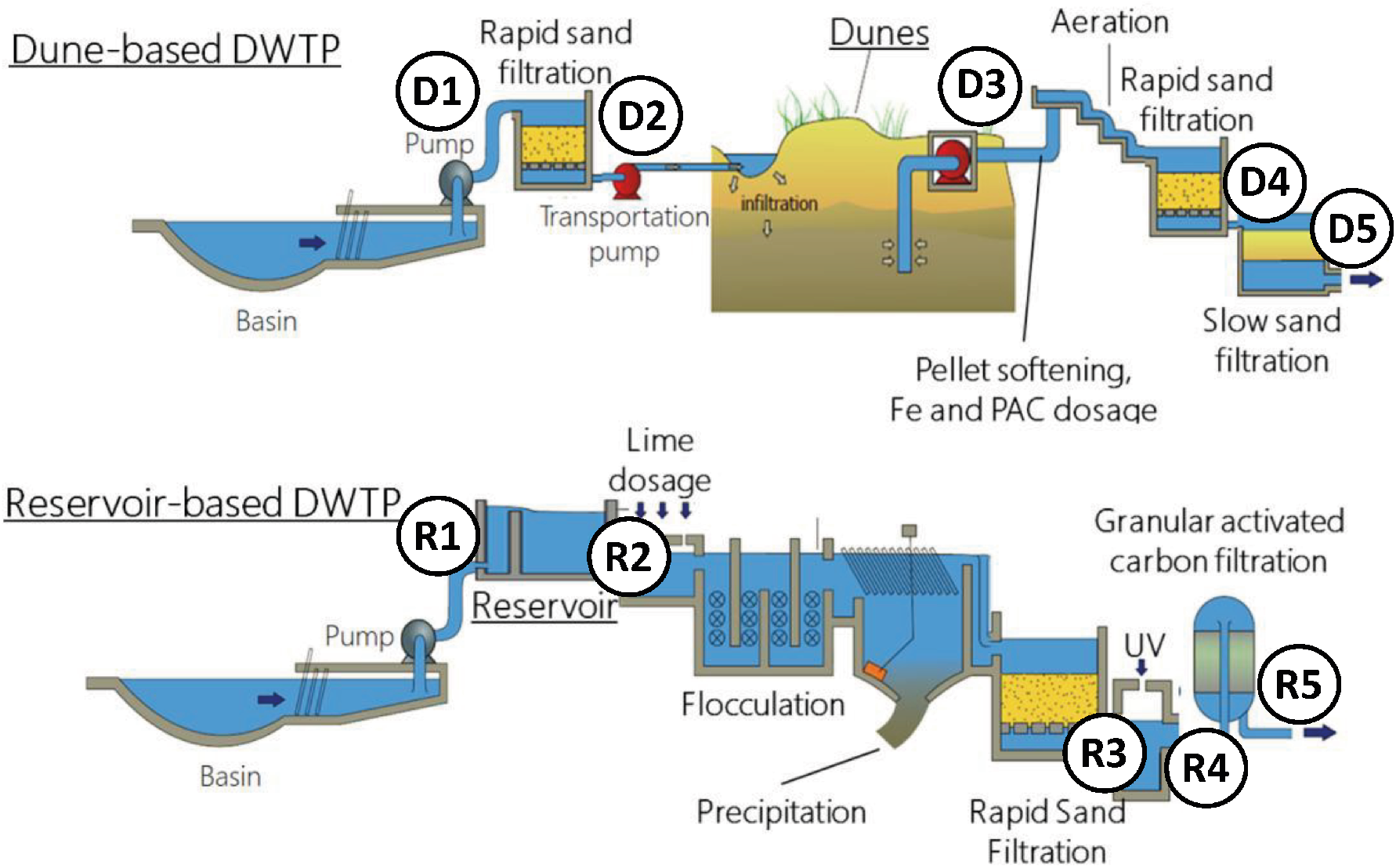
Schemes of the dune-based and reservoir-based drinking water treatment processes (DWTPs). The dunes and the reservoir are the storage water steps (underlined). The dune-based DWTP consists of a first rapid sand filtration (RSF1) of the Meuse River water followed by infiltration and storage in dunes (HRT = 3 months). Subsequently, the dune water is processed by pellet softening to regulate hardness, and iron (Fe) and powdered activated carbon (PAC) is dosed to improve color, odor and the performance of the second rapid sand filtration (RSF2). Finally, the water is disinfected via slow sand filtration (SSF). In the reservoir-based DWTP, the Meuse River water is stored in a reservoir (HRT = 3 months) followed by lime dosage to regulate hardness, flocculation, precipitation, and rapid sand filtration to reduce turbidity and disinfection with UV and taste and odor correction with granular activated carbon (GAC). The overall hydraulic residence times of the waters across the treatment trains amount to circa 3 months in both processes. Sampling points are represented by numbers D1-D5 for the dune-based DWTP (*top*) and R1-R5 for the reservoir-based DWTP (*bottom*). Figure adapted from (van Halem and Rietveld, 2014).

**Table 1.** Water samples were collected from the different stages of the dune-based and reservoir-based DWTPs processing surface water from the Meuse River and Meuse River, respectively, in the Netherlands. The metadata for the molecular biology analyses are given.

| DWTP system | Sample collection date | Sample description | Water volume extracted (L) | DNA extract quality (A260/A280, $\pm 0.1$) | DNA concentration in extract (ng/μL) | Metagenomic analyte identifier | BioSample IDs |
| --- | --- | --- | --- | --- | --- | --- | --- |
| Dune-based | 2020-10-26 | Meuse River influent | 5.8 | 1.8 | 71 | INF-DB-DWTP | SAMN31143551 |
|  | 2020-10-26 | RSF1 outlet | 9 | 1.8 | 27 | RSF1-DB-DWTP | SAMN31143552 |
|  | 2020-11-30 | After dune infiltration | 1200 | 1.8 | 30 | DUNE-DB-DWTP | SAMN31143553 |
|  | 2020-12-14 | RSF2 outlet | 1200 | 1.8 | 15.5 | RSF2-DB-DWTP | SAMN31143554 |
|  | 2020-12-16 | SSF outlet | 1200 | 1.8 | 98 | SSF-DB-DWTP | SAMN31143555 |
| Reservoir-based | 2020-12-19 | Meuse river influent | 1.5 | 1.8 | 34 | INF-RB-DWTP | SAMN31143556 |
|  | 2021-02-24 | Reservoir outlet | 1.5 | 1.8 | 10 | RES-RB-DWTP | SAMN31143557 |
|  | 2021-03-02 | Before UV | 100 | 1.8 | 11 | BefUV-RB-DWTP | SAMN31143558 |
|  | 2021-03-03 | After UV | 100 | 1.8 | 22 | AftUV-RB-DWTP | SAMN31143559 |
|  | 2021-03-05 | GAC outlet | 210 | 1.8 | 11 | GAC-RB-DWTP | SAMN31143560 |

The dune-based DWTP samples consisted of (D1) influent from the Meuse River water (N 51° 55’ 41.7288; E 4° 46’ 15.7404), (D2) outlet of the first rapid sand filtration, (D3) dune outlet (3 months hydraulic residence time), (D4) outlet of the second rapid sand filtration, and (D5) outlet of the slow sand filtration. The reservoir-based DWTP samples consisted of: (R1) influent of the reservoir from the Meuse River water (N 51° 45’ 39.3228; E 4° 46’ 8.6664), (R2) a sample of water after being stored in the reservoir for 3 months, (R3) rapid sand filtration treatment outlet, (R4) UV treatment outlet, and (R5) GAC outlet. Water quality parameters were provided by the DWTPs (Figure S2). The volume of each water sample depended on the expected biomass concentration at each stage, based on the author’s experience and knowledge of DWTP personnel.

### 2.2. DNA extraction

Each water sample was immediately filtered on the DWTP site through a 0.22 μm polyethersulfone membrane via vacuum filtration. The membrane containing the biological retentate was folded and introduced into the DNA extraction tubes. Total DNA was extracted using the DNeasy PowerWater DNA extraction kit (Qiagen, The Netherlands) following the manufacturers’ instructions. DNA qualities of the extracts were measured as absorbance ratio at 260 and 280 nm using a NanoDrop spectrophotometer. DNA concentrations were measured with a Qubit4 fluorometer (Thermo Fisher Scientific, USA). The DNA quality and concentration obtained for each sample is given in Table 1.

### 2.3. Library preparation, sequencing, quality control, and assembly

#### Preparation of metagenome libraries

The DNA analytes were sent to Novogene (Cambridge, United Kingdom) for metagenome library preparation and sequencing. A total amount of 1 μg DNA per sample was used as input material to prepare libraries that were generated using the NEBNext^®^ Ultra^™^ DNA Library Prep Kit for Illumina (NEB, USA) following manufacturer’s instructions; index codes were added to attribute sequences to each sample. In short, the DNA sample was fragmented by sonication into fragment sizes of 350 bp; the DNA fragments were end-polished, A-tailed, and ligated with the full-length adaptor for Illumina sequencing with further PCR amplification to add the sequence adapters; the PCR products were purified on AMPure XP magnetic beads (Beckman Coulter, USA).

#### Sequencing of libraries

The library preparations were sequenced with an Illumina HiSEQ PE150 system. Ten raw sequencing files with 150 bp paired-reads were obtained, with an average of 5.6 Gb per sample (41 million reads). More details are given in Table S1.

#### Quality control of sequenced reads

The quality of the sequenced raw reads was assessed by FastQC (version 0.11.7) with default parameters (Andrews, 2010) and visualized with MultiQC (version 1.0) (Figure S1). Low-quality paired-end reads were trimmed and filtered by Trimmomatic version 0.39 on the paired-end mode (Bolger et al., 2014).

#### Assembly of sequence reads

Clean reads were assembled into contigs using MetaSPAdes (version 3.13.0) with a contig length between 300 and 2000 bp (Nurk et al., 2017). The number of contigs obtained was 37,744.2 on average. Altogether, they had a total length of 180,840 kb, on average (Table S1).

### 2.4. Microbiome profiling obtained from dune and reservoir-based DWTPs

Taxonomic classification of raw reads was performed to profile the microbiome from each sample using the standard Kraken2 (version 2) database (uses all complete bacterial, archaeal, and viral genomes in NCBI Refseq database) with default parameters (Wood et al., 2019). Raw reads, divided into k-mers (substrings of length *k* contained within a biological sequence, determined by Kraken2), were matched with the NCBI database (Agarwala et al., 2018). The absolute abundance of each taxonomic group was indicated as the number of k-mers aligned to a specific taxonomic group. The relative abundance is the normalization of the total number of k-mers aligned in each sample.

Species richness (S) was measured as the number of different species detected in the raw datasets. The Shannon (H’) diversity index was calculated with the following equation:

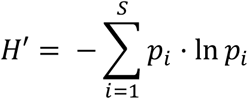

where *pi* represents the relative abundance of species *i* with respect to the total amount of species (S). Microbial community distance estimation was calculated using MinHash in Mash v2.3.(Ondov et al., 2016) with “-k” 18, the minimum value required for distance estimation.

### 2.5. Resistome and mobilome profiling of DNA analytes obtained from both DWTPs

ARGs were annotated by aligning the assembled contigs > 500 bp to the ResFinder 4.0 resistance gene database using the BLASTn (version 2.6.0) nucleotide alignment tool with a cut-off E-value <10^-5^ and sequence identity above 90% (Bortolaia et al., 2020). The richness of ARGs was defined as the number of different detected ARGs.

The mobilome was analyzed on the same set of contigs >500 bp using BLASTn (version 2.6.0) with the following specific databases of MGEs with sequence identity >95% and an e-value <10^-20^. The presence of plasmids was studied with the PLSDB database (Galata et al., 2019). Integrons were detected with the INTEGRALL database (Moura et al., 2009). The ISfinder database was used to identify bacterial insertion sequences (Siguier et al., 2006). The ICEberg database (version May 2, 2018) detected bacterial integrative and conjugative elements (Liu et al., 2019). For all queries, the ARG or MGE identified with the best score was selected to annotate the query.

Co-occurrence (or co-localization) of MGEs and ARGs within the same contig was identified. It was checked with the BLASTn outputs if a contig contained both ARGs and MGEs. Contigs >500 bp that simultaneously included hits from the ResFinder 4.0 database and at least one of the different MGE databases were considered to have co-localized. Afterward, a specific Kraken2 taxonomic analysis was performed with these contigs to identify the potential microbial host that might carry the co-localized ARG and MGE.

### 2.6. Functional analysis of antibiotic-producing microorganisms in water samples

Functional analysis of the metagenomes was performed to detect antibiotic synthesis pathways within microorganisms present in the microbial communities of the water samples. The assembled contigs were transcribed to coding sequences using Prokka (version 1.14.5) with default parameters (Seemann, 2014). Output files were introduced in GhostKOALA (version 2.2.) to assign protein functionality in the Kyoto Encyclopaedia of Genes and Genome s(KEGG) sequence library of *“genus_prokaryotes”* + *“family_eukaryotes”* (version 97.0). The mapping onto the KEGG pathway map was performed to obtain the antibiotic-resistance metabolic profile of the metagenome (Mitra et al., 2011).

### 2.7. High-throughput quantitative PCR analysis

Aliquots of the DNA extracts were sent in parallel to Resistomap (Helsinki, Finland) for high-throughput quantitative PCR (HT-qPCR) to detect and quantify the presence and abundance of 295 genes (listed in Table S2). These genes belonged to ARGs (238 genes), MGEs (51), and pathogens (6). A concentration of 2 ng DNA μL^-1^ in a reaction volume of 0.05 μL was used to obtain the number of gene copies of the different biomarkers. HT-qPCR results were corrected (detailed explanation in supplementary material) to get the number of gene copies existing per volume of filtered water from the DWTPs sampling points.

The gene abundance results were expressed in different ways. The absolute abundance of ARGs and MGEs was calculated as a number of gene copies per mL of filtered water as done in Xu et al. (2016). The absolute abundances of ARGs and MGEs sorted by antibiotic class and MGE type were averaged over all genetic components belonging to each group. The relative abundance of ARGs and MGEs was calculated based on the ARG or MGE copies per number of 16S rRNA gene copies. Abundance values were logarithmically transformed for comprehensive data calculation and visualization.

### 2.8. Statistics and data visualization

Graphs were made with RStudio (version 1.3.1093). Microbiome absolute and relative abundances were calculated by Pavian (Breitwieser and Salzberg, 2020). Linear correlations between absolute abundances of ARGs and MGEs were analyzed using Pearson correlation coefficient (“ggpubr” R package) at value < 0.05. Pearson correlations between ARGs and each specific type of MGE (plasmid, insertion sequence, integron, and transposon) were also calculated. This gives the first-hint proxy for examining the co-localization of ARGs and MGEs.

## 3. Results

### 3.1. Microbial community composition

#### 3.1.1. Richness and alpha diversity of the water metagenomes

The metagenomes of the microbial communities present at the different sampling points across the dune-based and reservoir-based DWTPs were sequenced to obtain first their taxonomic profiles. DNA extracted from 10 water samples was sequenced, resulting in 41 ± 5 million paired-end reads per sample (Table S1). All sequenced samples had high-quality rates (quality rate per sequence base > 30; Q = −10 x log10(P), where P is the probability that a base call is erroneous) (Figure S1). In analogy to engineered ecosystems, and likely owing to current databases incompleteness, an average of 24.6 ± 6.5 % of the raw reads were taxonomically classified.

The alpha diversity of the water microbiomes was assessed using the richness and the Shannon index (Figure 3A). The former measures the number of different populations (at the genus level) in the community, and the latter accounts for the number, relative abundance, and evenness of species (Hill et al., 2003). Richness was stable throughout both DWTPs, ranging between 7027 and 7959 different classified species detected from the water metagenomes (Figure 3). The richness of the influent of the reservoir-based DWTP was 7% higher than the dune-based DWTP influent water. Rapid sand filtration (RSF1 and RSF2), slow sand filtration (SSF), and granular activated carbon (GAC) increased the number of species by 2.9 ± 1.7%. On the contrary, dune infiltration, reservoir, and UV disinfection decreased it by 5.2 ± 2.1%. The Shannon H’ diversity index ranged between 4.9 and 7.7 across all samples (*i.e*., equivalent to 134 to 2208 virtual equi-abundant populations). The dune-based DWTP gradually increased from 5.3 to 7.7 throughout the plant. Equal H’ diversity values were found in the influent (6.7) and effluent (6.6) of the reservoir-based DWTP despite its oscillating trend.

**Figure 3.**
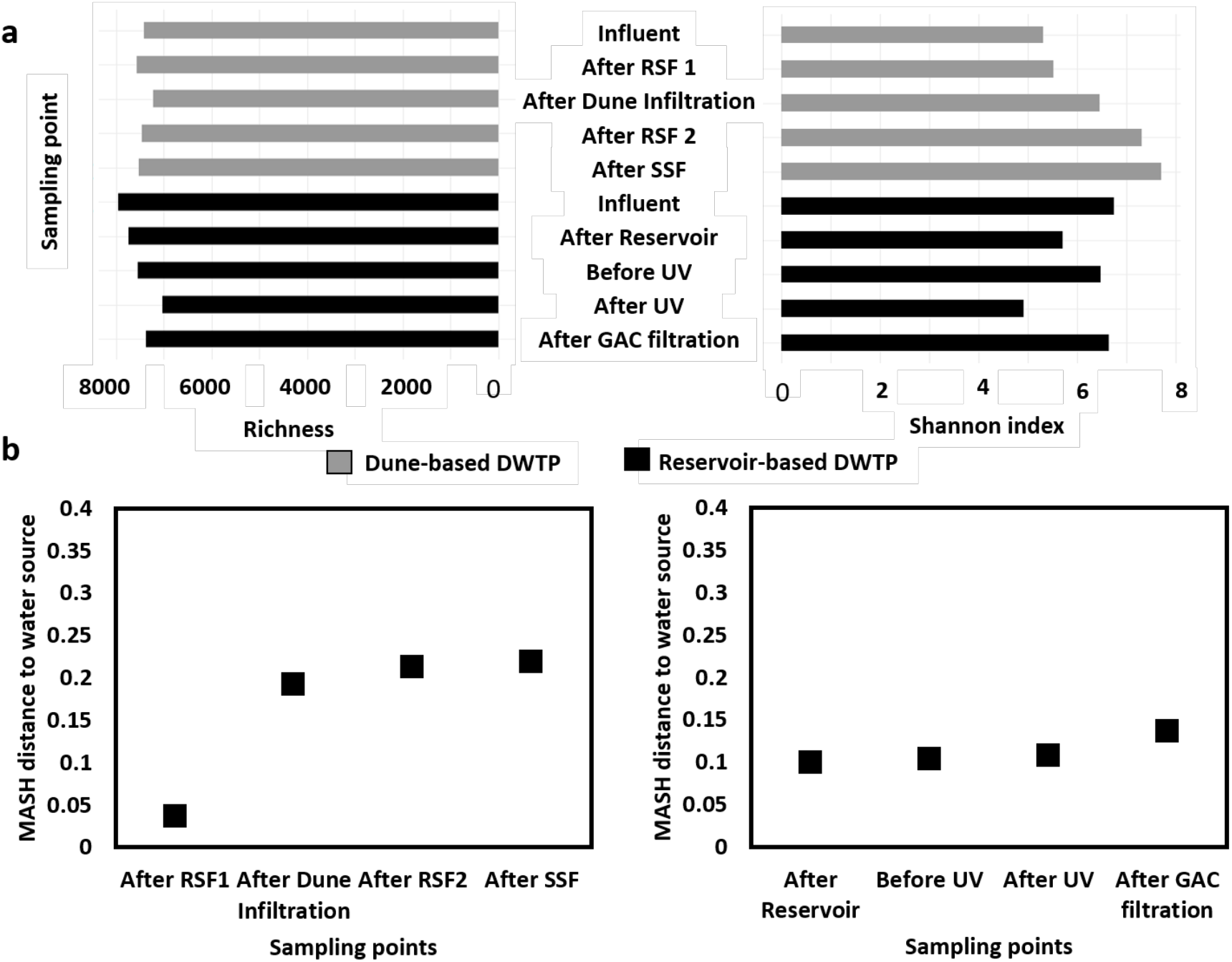
A) Richness and Shannon diversity indices (*x-axes*) of the taxonomically classified metagenomics datasets of the waters sampled at the different locations (*y-axes*) within the dune-based and reservoir-based DWTP trains. B) Microbial community distance estimation with MinHash between each sampling point and the influent (p-value < 0.05). RSF: rapid sand filtration; SSF: slow sand filtration; GAC: granular activated carbon filtration.

The distances between the microbial community compositions were calculated using the MinHash dimensionality-reduction technique in Mash (Figure 3B). Higher distance indicates the larger dissimilarity between the microbial community at each sampling point and the influent. The dissimilarity significantly increased after every step in both DWTPs (p-value < 0.05, expect for After Reservoir where p-value = 0.09). Overall, the differences in microbial community compositions across the process train of the dune-based DWTP were higher than in reservoir-based DWTP.

#### 3.1.2. Taxonomic classification of microbial communities

The relative abundance of the detected prokaryotic populations across the DWTPs at phylum and genus levels is shown in Figure 4. The river-influent water of both DWTPs had similar compositions. At phylum level, *Proteobacteria* (68.1 ± 5.5% in dune-based DWTP vs. 76.9 ± 10.2% in reservoir-based DWTP), *Actinobacteria* (17.1 ± 6.5% vs. 11.9 ± 5.6%) and *Bacteroidetes* (6.2 ± 3.8% vs. 4.9 ± 2.6%) dominated the microbial communities of both DWTPs. At genus level, the freshwater genera *Limnohabitans* (24.6% vs. 6.4%), *“Candidatus* Planktophila” (8.2% vs. 7.5%), and *Flavobacterium* (6.8% vs. 6.1%) were the main populations detected in both river waters. Their relative abundance decreased throughout the DWTP processes. Interestingly, we found several microbes in the effluent water that were absent in the influent, with most of them appearing after dune infiltration, SSF, and GAC filtration.

**Figure 4.**
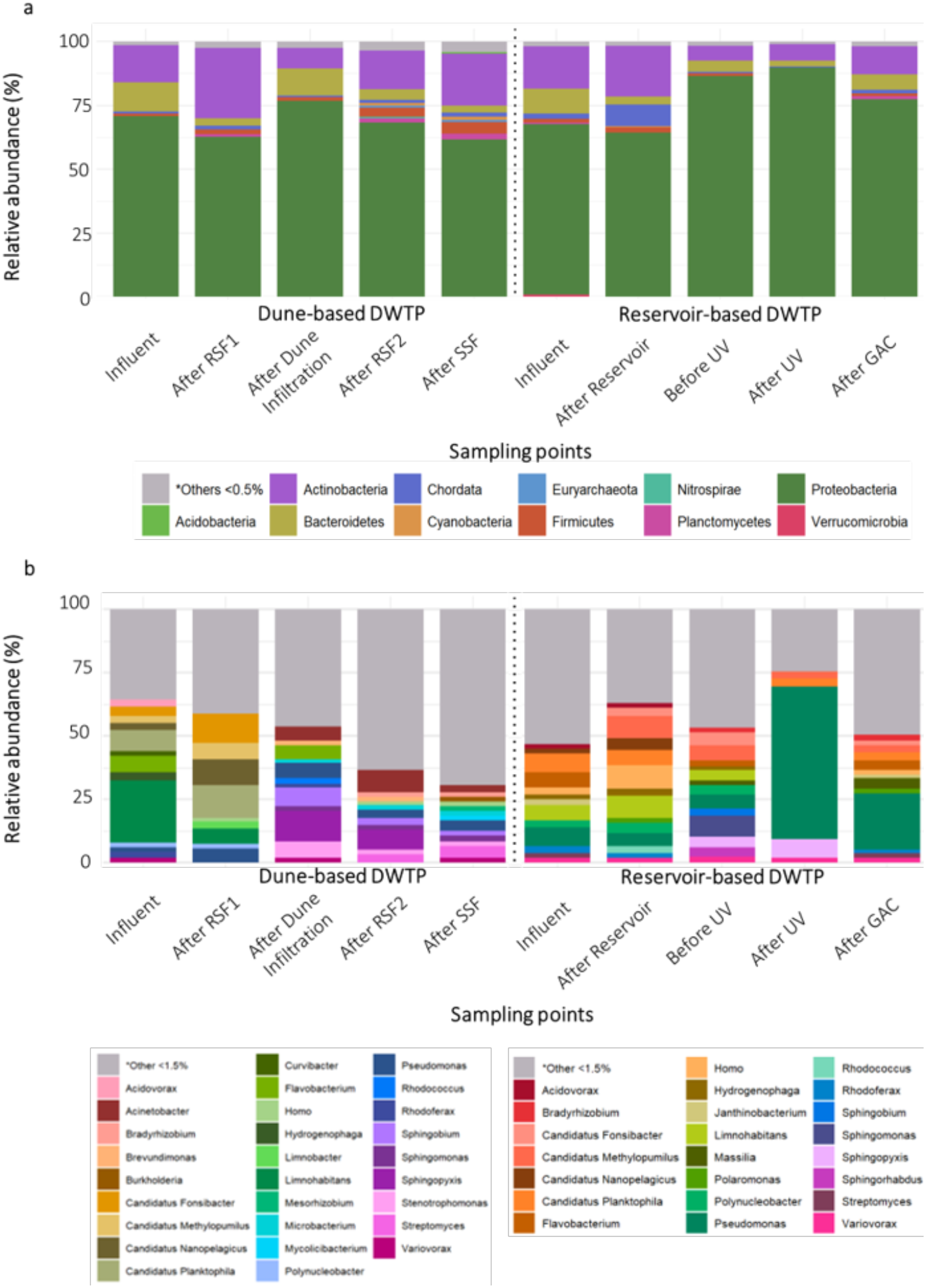
Microbial community composition at phylum (a) and genus (b) level of dune-based and reservoir-based DWTPs, as measured by metagenomics. The relative abundance of classified genera (Y-axis) is represented in the different sampling points (X-axis). Genera with less than 1.5% abundance in all samples were grouped as others.

In the dune-based DWTP, every sand filter decreased the relative abundance of members of the phylum *Bacteroidetes* (Figure S2) and its most abundant genus *Flavobacterium*. This population decreased from 6.8% to 0.8% in RSF1, from 5.4% to 1% in RSF2, and from 1% to 0.2% in SSF. In contrast, no other genus systematically increased after all sand filtration steps. The most notorious changes were the increase in the relative abundance of *Pseudomonas* (3.5%) and *Acinetobacter* (0.9%) in RSF1 and *Streptomyces* (4.5%) in SSF. Overall, the water infiltration in dunes had the highest impact on the microbial community composition: (i) it substantially decreased the relative abundance of genera that were abundant in the influent, namely *Limnohabitans* (from 6.1 to 0.2%) and of *“Ca*. Planktophila” (from 12.8 to <0.1%); and (ii) increased the relative abundances of other genera like *Sphingophyxis* (from 0.1 to 12.1%) and *Sphingobium* (from 0.2 to 12.7%).

Unlike dune infiltration in the dune-based DWTP, the water storage step in the reservoir-based DWTP did not drastically modify the microbial community of the water. In this DWTP, the most significant change took place in the disinfection step, UV disinfection. The relative abundance of *Pseudomonas* increased from 7.6 to 60%, and *Sphingopyxis* raised from 4.0 to 7.5%. Concomitantly, the presence of the other genera decreased. In the following unit operation, GAC filtration, the relative abundance of *Pseudomonas* decreased to 22.1%, whilst that of other genera such as *Massilia* (3.7%), *Polaromonas* (1.3%) and *Flavobacterium* (2.4%) increased.

### 3.2. Pathogenic bacteria decreased across both chlorine-free drinking water treatment plants

HT-qPCR was used to detect the presence of *Acinetobacter baumannii, Pseudomonas aeruginosa*, and Enterobacteriaceae, the three most critical antibiotic-resistant pathogenic bacteria as designated by the World Health Organization (Tacconelli and Magrini, 2017). Overall, the absolute abundance of the pathogenic bacteria detected by qPCR was low (< 10^6^ gene copies mL^-1^) and further reduced along the two DWTPs. *Acinetobacter baumanii, Pseudomonas aeruginosa*, and *Enterococci* were detected (>10^2^ genes copies mL^-1^), while *Klebsiella pneumoniae, Campylobacter*, and *Staphylococci* were not. *A. baumanii* was detected across both plants. *P. aeruginosa* was recalcitrant across the treatment train of the dune-based DWTP but was not detected after UV disinfection in the reservoir-based DWTP. Interestingly, *Enterococci* was found only after RSF1 in the dune-based DWTP.

### 3.3. Gram-negative bacteria as potential carriers of ARGs in DWTPs

The resistance determinants from the two DWTPs exhibited a large diversity of ARGs, highlighted by both qualitative (metagenomics) and quantitative (HT-qPCR) analyses.

The resistome richness ranged from 3 to 20 different ARGs detected per sample. In total, 34 different ARGs were detected in the water of the dune-based DWTP and 58 in the water of the reservoir-based DWTP (Figure 6). The most abundant ARGs related to resistance against macrolides (MLSB, 57 different ARGs), followed by beta-lactams (13), aminoglycosides (10), quinolones (7), sulfonamide (2), tetracycline (2), and trimethoprim (1). Some ARGs were recalcitrant across the DWTPs: notably, the *msr(D)_2_AF27302* gene conferring macrolide resistance remained in the treated water of the reservoir-based DWTP. In the dune-based DWTP, the dune infiltration step was most prominently increasing the diversity of ARGs, likely due to the drastic shift in microbial community (Figure 4). In the reservoir-based DWTP, the rapid sand filtration (before the UV step) and the GAC filtration introduced the highest variability in the resistome profile.

**Figure 5.**
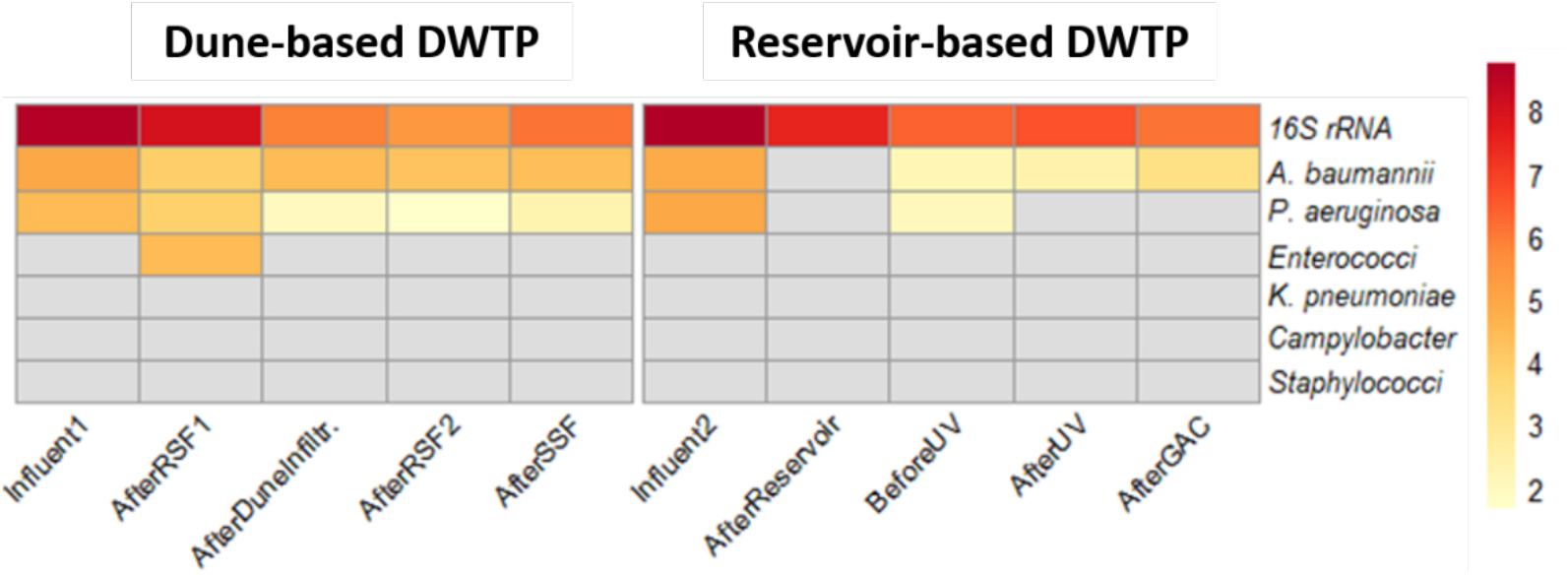
Heat map of absolute gene abundances (number of gene copies mL^-1^) of 6 pathogenic microorganisms in dune-based and reservoir-based DWTPs, displayed in logarithmic scale. Y-axis represents the abundance of the pathogens in the different sampling points of both DWTPs. In X-axis, the 16S rRNA absolute abundance is provided. RSF: rapid sand filtration; SSF: slow sand filtration; GAC: granular activated carbon filtration.

**Figure 6.**
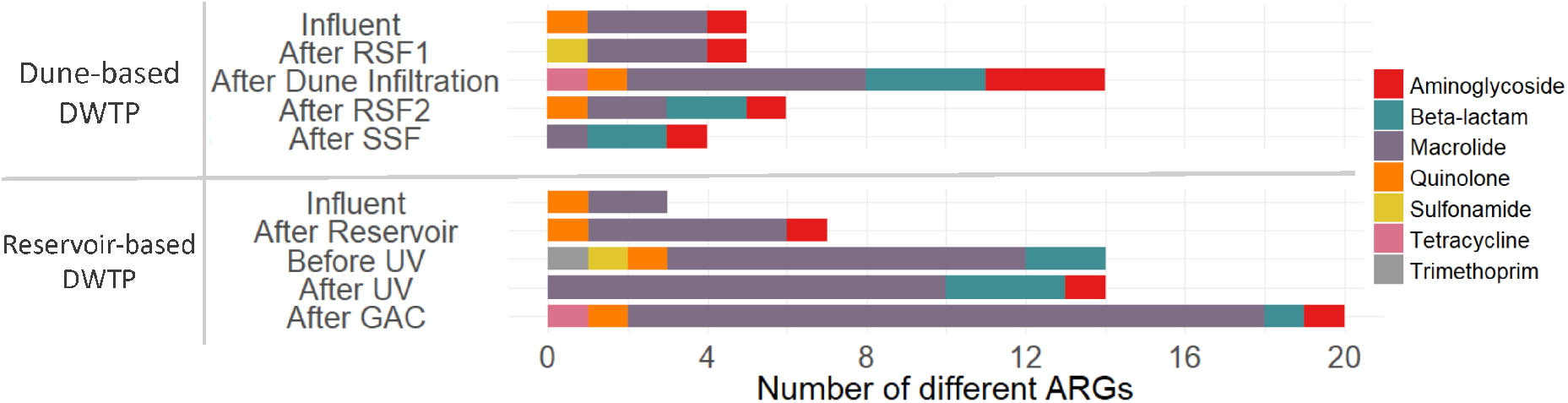
Resistome profile of dune-based and reservoir-based DWTP microbiome sorted by antibiotic class. The number of the different ARGs sorted per antibiotic class is represented in the different sampling points. The dotted line discriminates the results coming from each of the analyzed DWTPs.

We linked ARG contigs to potential microbial origins by assigning taxonomies to contigs carrying ARGs. The results of this analysis at genus level are given in Figure S3. Generally, contigs containing ARGs mainly affiliated with *Limnohabitants* in the influent water samples of both DWTPs. Other genera included *Paracoccus* and “*Ca*. Fonsibacter” in reservoir-based DWTP and *Polynucleobacter, Acidovorax, Hydrogenophaga*, and *“Ca*. Fonsibacter” in dunes-based DWTP. Most of these populations but “*Ca*. Fonsibacter” decreased across the treatment train in reservoir-based DWTP, while “*Ca*. Fonsibacter” and *Limnohabitans* persisted within the dune-based DWTP.

In the reservoir-based DWTP, the last GAC filtration step mostly increased the number of hosts carrying ARGs. This promoted the release of bacteria potentially carrying ARGs, such as *Pseudomonas, Kaistella, Microbacterium, Cellulosimicrobium, Caulobacter, Methylobacterium, Rhodoplanes, Messorzhibium*, and *Rhodoferax*, among others. In the dune-based DWTP, the dune infiltration step introduced the potential hosts carrying ARGs in the drinking water treatment train. *Acinetobacter, Rhodoferax*, and *Pseudomonas* were the microbial genera that persisted throughout the process after infiltration in the dune.

### 3.4. Chlorine-free DWTPs achieve 2-3 logs removal of ARGs and MGEs

HT-qPCR was used to assess the ARG and MGE removal efficiencies from both DWTPs by quantifying the number of gene copies per volume of filtered water in each sampling point. The absolute concentration of ARGs decreased along the treatment trains down to 2.2 log gene copies mL^-1^ (dune-based DWTP) and 2.6 log gene copies mL^-1^ (reservoir-based DWTP) (Figure 7a). MGEs decreased by 2.7 log gene copies mL^-1^ (dune-based DWTP) and 2.6 log gene copies mL^-1^ (reservoir-based DWTP) (Figure 7b). Similarly, the bacterial proxy *16S rRNA* gene decreased by 2.5 (dune-based DWTP) and 2.6 (reservoir-based DWTP) log gene copies mL^-1^.

**Figure 7.**
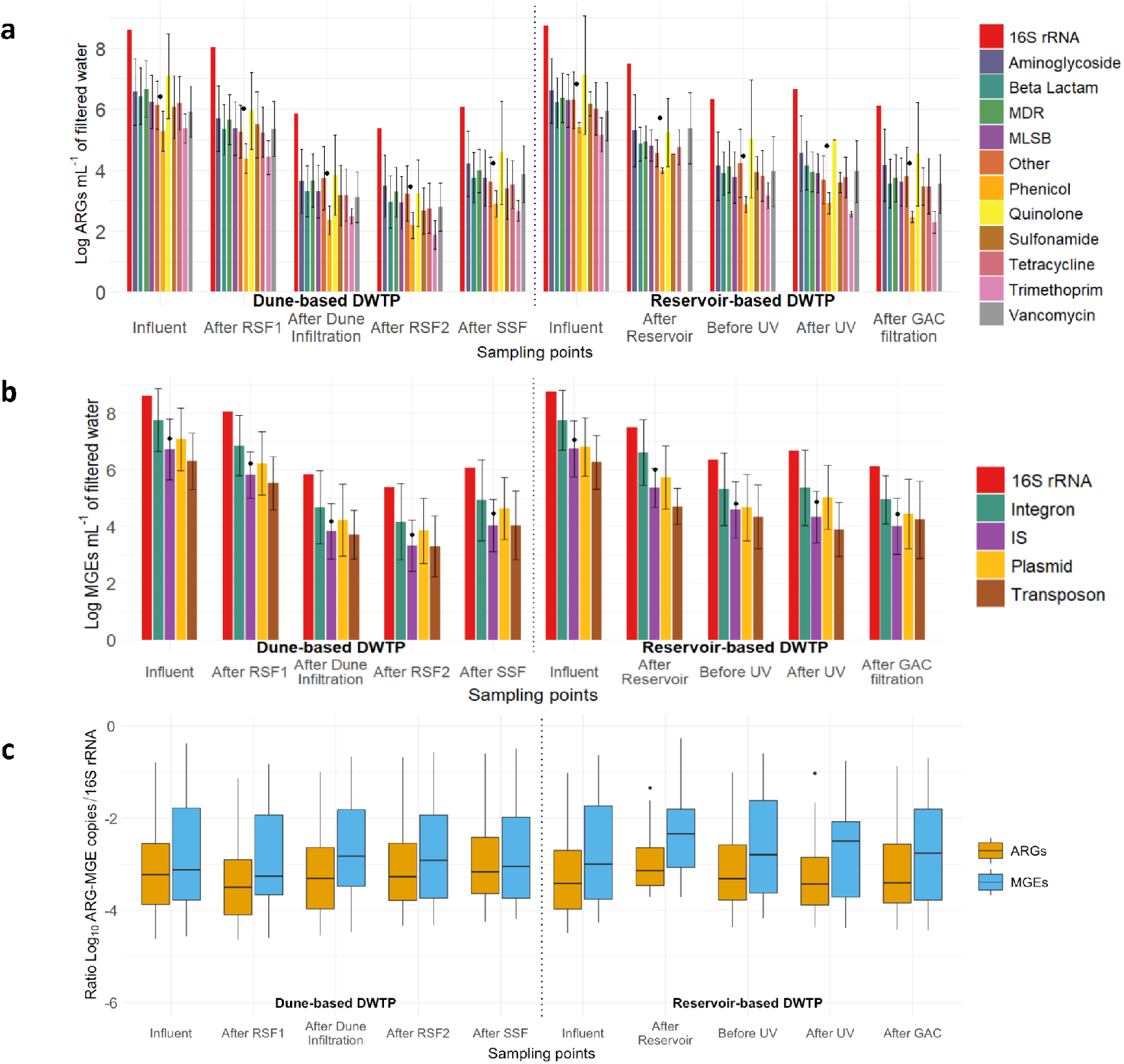
(a) Absolute abundance of difference ARGs mL^-1^ (sorted by antibiotic class) from both DWTPs. Values are represented on a logarithmic scale. Black dots indicate the average ARGs abundance per sampling point (b) Absolute abundance of difference MGEs mL^-1^ (sorted by antibiotic class) from both DWTPs. Values are represented on a logarithmic scale. Black dots indicate the average ARGs abundance per sampling point. (c) The ratio of ARGs or MGEs */16S rRNA* in both DWTPs. Each boxplot represents (from top to bottom) maximum, upper quartile, median, lower quartile, and minimum values. Note: RSF: rapid sand filtration; SSF: slow sand filtration; GAC: granular activated carbon.

The influent water samples from both DWTPs contained similar ARG concentration: 6.4 ± 0.9 (dune-based DWTP) and 6.8 ± 0.9 (reservoir-based DWTP) log ARG copies mL^-1^. Across the dune-based DWTP (Figure 7a), the concentration of ARGs evolved from 6.0 ± 0.9 (after the first rapid sand filtration) to 3.9 ± 0.9 (after dune infiltration), 3.5 ± 0.9 (after the second rapid sand filtration), and 4.2 ± 0.9 (after last slow sand filtration) log ARG copies mL^-1^. In the reservoir-based DWTP, the ARG concentration evolved from 5.7 ± 0.7 (after 3 months in reservoir) to 4.5 ± 0.9 (before UV treatment), 4.8 ± 0.8 (after UV treatment), and 4.2 ± 0.9 (after GAC filtration) log ARG copies mL^-1^.

The MGEs concentration in dune-based DWTP was 7.1 ± 1.2 log MGE copies mL^-1^ in the influent, 6.2 ± 1.1 log MGE copies mL^-1^ after RSF1 step, 4.2 ± 1.1 log MGE copies mL^-1^ after the dune infiltration, 3.7 ± 1.1 log MGE copies mL^-1^ after the RSF2 and 4.5 ± 1.1 log MGE copies mL^-1^ after the last SSF step. In the reservoir-based DWTP case, the MGE concentration in the influent was 7.1 ± 1.1 log MGE copies mL^-1^, followed by 6.0 ± 0.9 log MGE copies mL^-1^ after the 3 months’ time in the reservoir, 4.8 ± 1.2 log ARG copies mL^-1^ before UV treatment, 4.9 ± 1.1 log MGE copies mL^-1^ after UV treatment and 4.4 ± 1.2 log ARG copies mL^-1^ after GAC filtration (Figure 7b).

Some process stages increased the concentration of ARGs and MGEs in water, such as the slow sand filtration in the dune-based DWTP (22% in ARGs and 20% in MGEs) and the UV treatment in the reservoir-based DWTP (7% in ARGs and 2% in MGEs). However, the decrease in the concentration of ARGs and MGEs was progressive across both DWTPs. Detailed information on the ARGs and MGEs reduction throughout the processes per sampling point is given in Table S2.

Several ARGs persisted across both DWTPs. The *aadA7* (aminoglycoside resistance; 6.0 ± 1.2 log gene copies mL^-1^), *mexF* (multi-drug resistance; 5.6 ± 1.2) and *fox5* (beta-lactam resistance; and 5.5 ± 1.3 log gene copies mL^-1^) genes were the 3 most abundant ARGs in both DWTPs (Figure S4). Other ARGs were not present in the influent but appeared across the DWTPs such as *blaTEM, blaPAO* and *vanWG*. From the 238 ARGs tested, 72 *(i.e*., 30%) were not detected in any sampling point.

Regarding MGEs, the integron genes were the most abundant in both DWTPs (5.8 ± 1.7 log gene copies mL^-1^): the *intI1_1* (integron), *repA* (plasmid), *intI3* (integron), and *Tn5403* (transposon) genes were the most abundant (Figure S5). The conjugative plasmid sequences such as *IncP_oriT* and *trbC and* promiscuous plasmid *IncQ_oriT* gene sequences were also abundant.

Despite the reduction in ARGs and MGEs concentration in water, the ratios of ARGs and MGEs to the *16S rRNA* gene remained stable throughout both DWTPs (Figure 7c). This indicates that the DWTP process did not enrich for bacterial populations carrying ARGs or MGEs.

### 3.5. ARGs and MGEs co-localized on contigs of DWTP metagenomes

When ARGs and MGEs co-localize on the same genetic fragment, there is an increased chance that the fragment can be transferred between bacterial cells. Since facilitating the transfer, conjugation, integration, and transposition of genes in genomes, MGE can pose a risk for the dissemination of ARGs. Sets of 7 (dune-based DWTP) and 12 (reservoir-based DWTP) events of co-localization of ARGs and MGEs were detected on contigs retrieved from the sequenced metagenomes. A detailed description of the co-localization events is provided in Table S3. Co-localizations were detected in all sampling points from both DWTPs, except for the influent of the reservoir-based DWTP. The ARGs involved coded for mainly aminoglycosides and beta-lactam resistance.

For instance, the *ant(3”)-Ia* gene is an aminoglycoside resistant gene broadly described in *Klebsiella pneumoniae*. This persistent ARG was annotated from the metagenome of the dune-based DWTP influent and in the outlets of the RSF1, reservoir, UV, and GAC units. This ARG was embedded in plasmids, integrons, and bacterial integrative conjugative elements (ICEs), affiliating with *Polynucleobacter* and *Pseudomonas* genera. The *blaVIM* ARG against last-resource beta-lactamases (carbapenem antibiotics) was embedded in a plasmid, between insertion sequences, and in conjugative elements in different stages of the process: in the reservoir-based DWTP, it taxonomically affiliated with *Sphingobium* and *Sphingomonas* before UV, and with *Pseudomonas* after UV and GAC.

Other ARGs only appeared once. The *sul2* and *blaOXA-287* genes appeared after RSF1 and dune infiltration, respectively, both affiliating with *Acinetobacter*. The *sul2* gene was carried by plasmid, integrons, and bacterial integrative and conjugative elements. A plasmid carried the blaOXA-287 gene. The *sul1* gene appeared before UV, linked to *Sphingobium*. The *blaTEM-181* gene after UV linked to *Bacillus*. Both *sul1* and *blaTEM-181* genes were potentially carried by plasmids, integrons, and bacterial integrative and conjugative elements. The *srm(B)* gene after GAC was already assigned in *Rhodoferax*. The *mef(C)* and *mph(G)* genes appeared after GAC as well, with *Kaistella* as the potential host. These 2 ARGs were detected on the same contig (NODE_16870), which implied the possible existence of a plasmid co-containing multiple resistance genes.

## 4. Discussion

### 4.1. Operational units shape the structure of the drinking water microbiome

In order to study the impact of each unit operation on the dynamics of the water microbiome, we applied shotgun metagenomics on water samples collected along the treatment train of two chlorine-free DWTPs, namely a dune-based and a reservoir-based plant. The alpha diversities of both effluents, calculated as H’ Shannon diversity indices, were varied from 4.9 to 7.7 (average 6.3 ± 0.8). These values are comparable to other chlorine-free DWTPs (4.37 ± 0.1 – 6.02 ± 0.4; (Palomo et al., 2016), and significantly higher than the ones in plants with chemical disinfection (*ca*. 2 – 4) (Tiwari et al., 2021; Waak et al., 2019)(Dai et al., 2019). Noticeably, the alpha diversity increased after every biological filter (RSF1, RSF2, SSF, and GAC filtration), a likely consequence of direct seeding from biofilm detachment (Velten et al., 2011) (Lautenschlager et al., 2014).

*Proteobacteria, Actinobacteria*, and *Bacteroidetes* were the most abundant phyla in both DWTPs. This matched with previous observations (Lin et al., 2014; Oh et al., 2017; Su et al., 2018), and with the presence of *Actinobacteria* and *Bacteroidetes* in freshwater ecosystems (Neuenschwander et al., 2018; Warnecke et al., 2004). Interestingly, the microbial community after the first operational unit of both DWTPs, namely RSF in dune-based DWTP and reservoir in reservoir-based DWTP, was similar to the influent. However, the similarity decreased in the downstream stages of the DWTPs (Figure 3B). This aligned with the observations of Pinto et al. (2012), who highlighted that even though the source water seeds the drinking water microbiome, the unit operations shape the structure of the effluent microbial community.

The conditions within biological sand filters have different impacts on microorganisms fitness (Hu et al., 2020), yet Webster & Fierer (2019) postulated that changes in community composition before and after lab-scale biological sand filters are largely predictable. In this line, we found higher abundances of *Actinobacteria* the RSF and SSF effluents as compared to their influents in the dune-based DWTP (Figure 4a), in analogy to the high *Actinobacteria* abundance in bench-scale sand filters (Xu et al.(2020)). In contrast, the relative abundance of *Bacteroidetes* decreased after every biological sand filtration unit, similar to earlier reports (Mukherjee et al., 2016; Pfannes et al., 2015). Another example are the common freshwater bacteria *Limnohabitans, “Ca*. Planktophila”, “*Ca*. Nanopelagicus” and *Rhodoluna* (Hahn, 2016; Kasalický et al., 2013; Neuenschwander et al., 2018), abundant in the water influent but almost absent in the DWTPs effluents. Overall, our findings showcase common patterns in the effect of biological operational units unit on the water microbiome dynamics of full-scale DWTP, paving the way to predict and modulate the microbial community in the drinking water effluent.

### 4.2. Chlorine-free DWTPs remove antibiotic resistance determinants

To the best of our knowledge, this is the first study in which the fate of ARG and ARB is monitored throughout the treatment train of chlorine-free DWTPs. Both DWTPs effectively reduced the concentration of ARGs by *ca*. 2.5 log gene copies mL^-1^. These removals are comparable to the highest reported in chlorine-amended DWTPs, between <0.1 log ARG copies mL^-1^ (Su et al., 2018; Zhang et al., 2016) and 2.4 log ARG copies mL^-1^ (Hu et al., 2019). Moreover, the total ARG concentration in the water effluent of both chlorine-free DWTPs was *ca*. 4 log ARG copies mL^-1^, similar to what Hu et al. (2019) found in a chlorine-amended DWTPs. Overall, both chlorine-free DWTP proved at least as effective as chlorine-amended DWTP at reducing ARGs. Additionally, the decrase in ARGs and MGEs concentration was linearly correlated with that of *16S rRNA* (Figure 7c, Figure S6), proving that none of the biological unit operations in chlorine-free DWTPs selected for ARB, *i.e*. the ARB/16S ratio did not increase.

The water storage steps yielded the highest ARG and ARB removal in both plants (2.1 and 1.6 ARG copies mL^-1^ in dune- and reservoir-based DWTPs, respectively), likely due to biomass decay and plasmid degradation due to the high hydraulic retention times and low nutrient availability in these systems (Amarasiri et al., 2020; Griffiths et al., 1990; Zhang et al., 2021). Likewise, RSFs reduced the ARG and ARB concentration by decreasing biomass concentration, as previously reported (Hu et al., 2019; Su et al., 2018; Zhang et al., 2016). Unexpectedly, GAC filtration also decreased the concentration of ARGs (Figure 7a), in contrast to previous studies describing GAC filtration as the critical step where resistance determinants increase (Su et al., 2018; Wan et al., 2021; Zhang et al., 2019). However, the decrease in ARGs concentration in this study contrasted with the increase in ARG richness (Figure 6), which suggests that the microbiome in the GAC effluent is seeded by the GAC biofilm.

The final treatment step before discharge to the environment is disinfection, which is intended to suppress or inactivate harmful microorganisms and prevent the regrowth of opportunistic bacteria (National Research Council Safe Drinking Water Committee, 1980). However, the SSF (0.75 log gene copies mL^-1^) and UV treatments (0.32 log gene copies mL^-1^) in this study increased the concentration ARGs (Figure 7a-b). The fate of ARG in SSF has not been directly studied before. However, Xu et al. (2020) showed that SSF hardly decreases the concentration of antibiotics in water, and Ciric (2022) reported that while their SSF removed most of the microorganisms, those in the effluent were more prone to resistance to antibiotics. In the case of UV treatment, previous studies proved its efficacy for cell reduction (plate counting) but not for ARGs removal (Chen and Zhang, 2013; Stange et al., 2019). Gram-negative bacteria, and specifically *Pseudomonas*, tolerate UV by efficient repair mechanisms, high growth rates, or the use of low-molecular-weight organic carbon (generated by UV illumination) as an energy source (Chen et al., 2020). This can explain the drastic rise in relative abundance of *Pseudomonas*, a common multi-drug resistant bacteria in drinking water systems (Su et al., 2018), and the quantitative increase in the *16S rRNA* gene marker after UV disinfection (Figure 7a). Nevertheless, despite the intermediate increase of ARGs, MGEs, and pathogenic bacteria after disinfection, DWTPs successfully reduced their effluent concentration (Figure 5, Figure 7, Figure S7).

### 4.3. Clinical implications of gene transfer in chlorine-free DWTPs

In concert with wastewater treatment plants (WWTPs), DWTPs are the ultimate barriers preventing the spread of waterborne diseases, and the release of antibiotics, ARB, and ARGs into water systems (Collignon and McEwen, 2019; Miłobedzka et al., 2022). One crucial aspect is the presence of ARGs against last-resort antibiotics such as carbapenems or colistin. Carbapenem resistance genes like *blaIMP or blaVIM* (class B beta-lactamases resistant genes) have been described in pathogenic bacteria such as *Pseudomonas, Acinetobacter*, or *Enterobacteriaceae* (Nordmann et al., 2012; Shanthi Amudhan et al., 2012). Carbapenem is a beta-lactam antibiotic with a broad antimicrobial spectrum and administered as a last resort for treating drug-resistant bacterial infections. However, the number of carbapenem-resistant bacteria has steadily increased (WHO, 2017), and represents a primary concern in drinking water. *blaIMP* genes were rarely detected along both DWTPs. However, *blaVIM* was detected along the dune-based DWTP and in the effluent (after GAC treatment) of the reservoir-based DWTP (Figure S4). The taxonomic annotation of the contigs containing *blaVIM* genes revealed their potential co-localization with multiple plasmids affiliating with *Sphingobium* in the dune-based DWTP, and with *Sphingobium* (plant pathogen) and *Pseudomonas* in the reservoir-based DWTP after GAC filtration. The carbapenem-resistant *Pseudomonas* is accounted by WHO within the list of critical priority pathogens for which new antibiotics are required (Tacconelli and Magrini, 2017). Colistin resistance genes, such as *mcr1* variants, were also highly abundant in both the dune-based and the reservoir-based DWTPs. However, this is not unique to chlorine-free DWTPs as multiple last-resort ARGs have also been identified in conventional DWTP (with chlorine use) as well as in tap water (Dias et al., 2020). Importantly, the *mcr1* gene load decreased significantly along the water treatment trains and neither co-localized with any MGE nor affiliated with any pathogenic bacteria. Further research should underpin the regrowth capacity of such pathogens in chlorine-free DWTP effluents.

## 5. Conclusions

In this work, we characterized for the very first time the abundance and dynamics of microbial communities, antibiotic resistance genes (ARGs), and mobile genetic elements (MGEs) across the treatment trains of two chlorine-free drinking water treatment plants. The in depth analysis of the metagenomes and resistomes led to the following main conclusions:

1. Chlorine-free DWTPs do not select for antibiotic resistant bacteria, as supported by the linear correlated between ARGs and MGEs, and the *16S rRNA* concentrations.
2. The measured reduction in ARGs concentration by *ca*. 2.5 log gene copies mL^-1^ in both chlorine-free DWTPs is comparable to the highest removals reported so far for chlorine-amended DWTPs.
3. Water storage systems alone reduced the abundance of the *16S rRNA* gene, ARGs, and MGEs by *ca*. 1.6 log gene copies mL^-1^, and dune infiltration achieved the highest removal.
4. Despite a *ca*. 2.5 log *16S rRNA* gene copies mL^-1^ reduction, the effluent microbial diversity increased likely due to the seeding from the biofilms actively growing in the rapid and slow sand filters, and the granular activated carbon ones.
5. Despite the overall ARG decrease in the DWTP, disinfection (slow sand filtration and UV radiation) internally increased the concentration of ARGs, MGEs, and 16S rRNA genes by *ca*. 0.5 log gene copies mL^-1^, yet with no impact on overall reduction.

Overall, our findings confirm the effectiveness of chlorine-free DWTPs in providing safe drinking water and reducing the load of antibiotic resistance determinants, offering the Water Authorities the possibility to establish centralized risk management around these specific treatment steps.

## 6. Data availability

Metagenome sequencing data were deposited in the NCBI database with the BioProject ID: PRJNA886863.

## 7. Authors’ contributions

DCF and FCR designed the study with inputs of MvL, DvH, ML, and DGW. Fieldwork was supported by BP and DdR. DCF, FCR, and MCS performed the experimental investigations. DCF and FCR wrote the manuscript with direct contribution, edits, and critical feedback by all authors. DCF and FCR are first co-authors of this manuscript.

## 8. Acknowledgements

This work is part of the research project “Transmission of Antimicrobial Resistance Genes and Engineered DNA from Transgenic Biosystems in Nature” (TARGETBIO) funded by the programme Biotechnology & Safety of the Ministry of Infrastructure and Water Management (grant no. 15812) of the Applied and Engineering Sciences (TTW) Division of the Dutch Research Council (NWO) (PhD thesis of David Calderón Franco). The work was further supported by the partnership program Dunea–Vitens: Sand Filtration partnership (17830) of the Dutch Research Council (NWO) and the drinking water companies Vitens and Dunea (PhD thesis of Francesc Corbera Rubio). Michele Laureni was supported by a VENI grant from the Dutch Research Council (NWO) (project number VI.Veni.192.252). Marcos Cuesta Sanz benefitted from an Erasmus+ grant.

## References

Amarasiri, M., Sano, D., Suzuki, S., 2020. Understanding human health risks caused by antibiotic resistant bacteria (ARB) and antibiotic resistance genes (ARG) in water environments: Current knowledge and questions to be answered. Crit. Rev. Environ. Sci. Technol. 50, 2016–2059. https://doi.org/10.1080/10643389.2019.1692611

Balcázar, J.L., Subirats, J., Borrego, C.M., 2015. The role of biofilms as environmental reservoirs of antibiotic resistance. Front. Microbiol. 6, 1216. https://doi.org/10.3389/FMICB.2015.01216/BIBTEX

Bolger, A.M., Lohse, M., Usadel, B., 2014. Trimmomatic: A flexible trimmer for Illumina sequence data. Bioinformatics 30, 2114–2120. https://doi.org/10.1093/bioinformatics/btu170

Bortolaia, V., Kaas, R.S., Ruppe, E., Roberts, M.C., Schwarz, S., Cattoir, V., Philippon, A., Allesoe, R.L., Rebelo, A.R., Florensa, A.F., Fagelhauer, L., Chakraborty, T., Neumann, B., Werner, G., Bender, J.K., Stingl, K., Nguyen, M., Coppens, J., Xavier, B.B., Malhotra-Kumar, S., Westh, H., Pinholt, M., Anjum, M.F., Duggett, N.A., Kempf, I., Nykäsenoja, S., Olkkola, S., Wieczorek, K., Amaro, A., Clemente, L., Mossong, J., Losch, S., Ragimbeau, C., Lund, O., Aarestrup, F.M., 2020. ResFinder 4.0 for predictions of phenotypes from genotypes. J. Antimicrob. Chemother. 75. https://doi.org/10.1093/jac/dkaa345

Breitwieser, F.P., Salzberg, S.L., 2020. Pavian: Interactive analysis of metagenomics data for microbiome studies and pathogen identification. Bioinformatics 36, 1303–1304. https://doi.org/10.1093/bioinformatics/btz715

Calderón-Franco, D., Sarelse, R., Christou, S., Pronk, M., van Loosdrecht, M.C.M., Abeel, T., Weissbrodt, D.G., 2022. Metagenomic profiling and transfer dynamics of antibiotic resistance determinants in a full-scale granular sludge wastewater treatment plant. Water Res. https://doi.org/10.1016/j.watres.2022.118571

Chen, H., Zhang, M., 2013. Effects of advanced treatment systems on the removal of antibiotic resistance genes in wastewater treatment plants from Hangzhou, China. Environ. Sci. Technol. 47, 8157–8163. https://doi.org/10.1021/es401091y

Chen, P.F., Zhang, R.J., Huang, S. Bin, Shao, J.H., Cui, B., Du, Z.L., Xue, L., Zhou, N., Hou, B., Lin, C., 2020. UV dose effects on the revival characteristics of microorganisms in darkness after UV disinfection: Evidence from a pilot study. Sci. Total Environ. 713, 136582. https://doi.org/10.1016/j.scitotenv.2020.136582

Ciric, L., 2022. ARE VICTORIAN WATER TREATMENT TECHNOLOGIES FIT FOR THE AMR ERA? [WWW Document]. Microbiol. Soc. URL https://microbiologysociety.org/our-work/75th-anniversary-a-sustainable-future/antimicrobial-resistance-amr/antimicrobial-resistance-amr-case-studies/victorian-water-treatment-technologies-amr.html

Clifford, R.J., Milillo, M., Prestwood, J., Quintero, R., Zurawski, D. V., Kwak, Y.I., Waterman, P.E., Lesho, E.P., Mc Gann, P., 2012. Detection of bacterial 16S rRNA and identification of four clinically important bacteria by real-time PCR. PLoS One 7, 7–12. https://doi.org/10.1371/journal.pone.0048558

Collignon, P.J., McEwen, S.A., 2019. One health-its importance in helping to better control antimicrobial resistance. Trop. Med. Infect. Dis. 4. https://doi.org/10.3390/tropicalmed4010022

Dai, Z., Sevillano-Rivera, M.C., Calus, S.T., Melina Bautista-de los Santos, Q., Murat Eren, A., van der Wielen, P.W.J.J., Ijaz, U.Z., Pinto, A.J., 2019. Disinfection exhibits systematic impacts on the drinking water microbiome. bioRxiv 1–19. https://doi.org/10.1101/828970

Destiani, R., Templeton, M.R., 2019. Chlorination and ultraviolet disinfection of antibiotic-resistant bacteria and antibiotic resistance genes in drinking water. AIMS Environ. Sci. 6, 222–241. https://doi.org/10.3934/environsci.2019.3.222

Dias, M.F., da Rocha Fernandes, G., Cristina de Paiva, M., Christina de Matos Salim, A., Santos, A.B., Amaral Nascimento, A.M., 2020. Exploring the resistome, virulome and microbiome of drinking water in environmental and clinical settings. Water Res. 174, 115630. https://doi.org/10.1016/j.watres.2020.115630

Farkas, A., Butiuc-Keul, A., Ciatarâş, D., Neamţu, C., Crăciunaş, C., Podar, D., Drăgan-Bularda, M., 2013. Microbiological contamination and resistance genes in biofilms occurring during the drinking water treatment process. Sci. Total Environ. 443, 932–938. https://doi.org/10.1016/j.scitotenv.2012.11.068

Galata, V., Fehlmann, T., Backes, C., Keller, A., 2019. PLSDB: A resource of complete bacterial plasmids. Nucleic Acids Res. 47, D195–D202. https://doi.org/10.1093/nar/gky1050

Griffiths, R.P., Moyer, C.L., Caldwell, B.A., Ye, C., Morita, R.Y., 1990. Long-term starvation-induced loss of antibiotic resistance in bacteria. Microb. Ecol. 19, 251–257. https://doi.org/10.1007/BF02017169

He, J.W., Jiang, S., 2009. Quantification of enterococci and human adenoviruses in environmental samples by real-time PCR (Applied and Environmental Microbiology (2005) 71:5 (2250-2255)). Appl. Environ. Microbiol. 75, 557. https://doi.org/10.1128/AEM.02602-08

Hill, T.C.J., Walsh, K.A., Harris, J.A., Moffett, B.F., 2003. Using ecological diversity measures with bacterial communities. FEMS Microbiol. Ecol. 43, 1–11. https://doi.org/10.1016/S0168-6496(02)00449-X

Hu, Y., Zhang, T., Jiang, L., Luo, Y., Yao, S., Zhang, D., Lin, K., Cui, C., 2019. Occurrence and reduction of antibiotic resistance genes in conventional and advanced drinking water treatment processes. Sci. Total Environ. 669, 777–784. https://doi.org/10.1016/j.scitotenv.2019.03.143

Huang, Z., Zhao, W., Xu, T., Zheng, B., Yin, D., 2019. Occurrence and distribution of antibiotic resistance genes in the water and sediments of Qingcaosha Reservoir, Shanghai, China. Environ. Sci. Eur. 31. https://doi.org/10.1186/s12302-019-0265-2

Jia, S., Bian, K., Shi, P., Ye, L., Liu, C.H., 2020. Metagenomic profiling of antibiotic resistance genes and their associations with bacterial community during multiple disinfection regimes in a full-scale drinking water treatment plant. Water Res. 176, 115721. https://doi.org/10.1016/j.watres.2020.115721

Jia, S., Shi, P., Hu, Q., Li, B., Zhang, T., Zhang, X.X., 2015. Bacterial Community Shift Drives Antibiotic Resistance Promotion during Drinking Water Chlorination. Environ. Sci. Technol. 49, 12271–12279. https://doi.org/10.1021/acs.est.5b03521

Khatri, N., Tyagi, S., 2015. Influences of natural and anthropogenic factors on surface and groundwater quality in rural and urban areas. Front. Life Sci. 8, 23–39. https://doi.org/10.1080/21553769.2014.933716

Kumpel, E., Cock-Esteb, A., Duret, M., Waal, O. De, Khush, R., 2017. Seasonal variation in drinking and domestic water sources and quality in port harcourt, Nigeria. Am. J. Trop. Med. Hyg. 96, 437–445. https://doi.org/10.4269/ajtmh.16-0175

Lautenschlager, K., Hwang, C., Ling, F., Liu, W.T., Boon, N., Köster, O., Egli, T., Hammes, F., 2014. Abundance and composition of indigenous bacterial communities in a multi-step biofiltration-based drinking water treatment plant. Water Res. 62, 40–52. https://doi.org/10.1016/j.watres.2014.05.035

Lin, W., Yu, Z., Zhang, H., Thompson, I.P., 2014. Diversity and dynamics of microbial communities at each step of treatment plant for potable water generation. Water Res. 52, 218–230. https://doi.org/10.1016/j.watres.2013.10.071

Liu, M., Li, X., Xie, Y., Bi, D., Sun, J., Li, J., Tai, C., Deng, Z., Ou, H.Y., 2019. ICEberg 2.0: An updated database of bacterial integrative and conjugative elements. Nucleic Acids Res. 47, D660–D665. https://doi.org/10.1093/nar/gky1123

Lübeck, P.S., Wolffs, P., On, S.L.W., Ahrens, P., Rådström, P., Hoorfar, J., 2003. Toward an international standard for PCR-based detection of food-borne thermotolerant campylobacters: Assay development and analytical validation. Appl. Environ. Microbiol. 69, 5664–5669. https://doi.org/10.1128/AEM.69.9.5664-5669.2003

Ma, L., Li, B., Jiang, X.T., Wang, Y.L., Xia, Y., Li, A.D., Zhang, T., 2017. Catalogue of antibiotic resistome and host-tracking in drinking water deciphered by a large scale survey. Microbiome 5, 154. https://doi.org/10.1186/s40168-017-0369-0

McConnell, M.M., Truelstrup Hansen, L., Jamieson, R.C., Neudorf, K.D., Yost, C.K., Tong, A., 2018. Removal of antibiotic resistance genes in two tertiary level municipal wastewater treatment plants. Sci. Total Environ. 643, 292–300. https://doi.org/10.1016/j.scitotenv.2018.06.212

Miłobedzka, A., Ferreira, C., Vaz-Moreira, I., Calderón-Franco, D., Gorecki, A., Purkrtova, S., Jan Bartacek, Dziewit, L., Singleton, C.M., Nielsen, P.H., Weissbrodt, D.G., Manaia, C.M., 2022. Monitoring antibiotic resistance genes in wastewater environments: The challenges of filling a gap in the One-Health cycle. J. Hazard. Mater. 424. https://doi.org/10.1016/j.jhazmat.2021.127407

Mitra, S., Rupek, P., Richter, D.C., Urich, T., Gilbert, J.A., Meyer, F., Wilke, A., Huson, D.H., 2011. Functional analysis of metagenomes and metatranscriptomes using SEED and KEGG. BMC Bioinformatics 12, 1–8. https://doi.org/10.1186/1471-2105-12-S1-S21

Mouchet, P., 1992. From conventional to biological removal of iron and manganese in France. J. / Am. Water Work. Assoc. 84, 158–167. https://doi.org/10.1002/j.1551-8833.1992.tb07342.x

Moura, A., Soares, M., Pereira, C., Leitão, N., Henriques, I., Correia, A., 2009. INTEGRALL: A database and search engine for integrons, integrases and gene cassettes. Bioinformatics 25, 1096–1098. https://doi.org/10.1093/bioinformatics/btp105

Mukherjee, N., Bartelli, D., Patra, C., Chauhan, B. V., Dowd, S.E., Banerjee, P., 2016. Microbial diversity of source and point-of-use water in rural Haiti - A pyrosequencing-based metagenomic survey. PLoS One 11, e0167353. https://doi.org/10.1371/journal.pone.0167353

National Research Council (US) Safe Drinking Water Committee, 1980. Drinking Water and Health, in: Drinking Water and Health: Volume 2. National Academies Press (US), pp. 955–956. https://doi.org/10.1111/j.1752-1688.1984.tb04810.x

Neuenschwander, S.M., Ghai, R., Pernthaler, J., Salcher, M.M., 2018. Microdiversification in genome-streamlined ubiquitous freshwater Actinobacteria. ISME J. 12, 185–198. https://doi.org/10.1038/ismej.2017.156

Nordmann, P., Dortet, L., Poirel, L., 2012. Carbapenem resistance in Enterobacteriaceae: Here is the storm! Trends Mol. Med. 18, 263–272. https://doi.org/10.1016/j.molmed.2012.03.003

Oh, S., Hammes, F., Liu, W.T., 2017. Metagenomic characterization of biofilter microbial communities in a full-scale drinking water treatment plant. Water Res. 128, 278–285. https://doi.org/10.1016/j.watres.2017.10.054

Ondov, B.D., Treangen, T.J., Melsted, P., Mallonee, A.B., Bergman, N.H., Koren, S., Phillippy, A.M., 2016. Mash: Fast genome and metagenome distance estimation using MinHash. Genome Biol. 17, 1–14. https://doi.org/10.1186/S13059-016-0997-X/FIGURES/5

Pallares-Vega, R., Blaak, H., van der Plaats, R., de Roda Husman, A.M., Hernandez Leal, L., van Loosdrecht, M.C.M., Weissbrodt, D.G., Schmitt, H., 2019. Determinants of presence and removal of antibiotic resistance genes during WWTP treatment: A cross-sectional study. Water Res. 161, 319–328. https://doi.org/10.1016/j.watres.2019.05.100

Palomo, A., Jane Fowler, S., Gülay, A., Rasmussen, S., Sicheritz-Ponten, T., Smets, B.F., 2016. Metagenomic analysis of rapid gravity sand filter microbial communities suggests novel physiology of Nitrospira spp. ISME J. 10, 2569–2581. https://doi.org/10.1038/ismej.2016.63

Pfannes, K.R., Langenbach, K.M.W., Pilloni, G., Stührmann, T., Euringer, K., Lueders, T., Neu, T.R., Müller, J.A., Kästner, M., Meckenstock, R.U., 2015. Selective elimination of bacterial faecal indicators in the Schmutzdecke of slow sand filtration columns. Appl. Microbiol. Biotechnol. 99, 10323–10332. https://doi.org/10.1007/s00253-015-6882-9

Pinto, A.J., Xi, C., Raskin, L., 2012. Bacterial community structure in the drinking water microbiome is governed by filtration processes. Environ. Sci. Technol. 46, 8851–8859. https://doi.org/10.1021/es302042t

Rook, J.J., 1976. Haloforms in Drinking Water. J. Am. Water Works Assoc. 68, 168–172.https://doi.org/10.1002/J.1551-8833.1976.TB02376.X

Sedlak, D.L., Von Gunten, U., 2011. The chlorine dilemma. Science (80-.). https://doi.org/10.1126/science.1196397

Seemann, T., 2014. Prokka: Rapid prokaryotic genome annotation. Bioinformatics 30, 2068–2069. https://doi.org/10.1093/bioinformatics/btu153

Sevillano, M., Dai, Z., Calus, S., Bautista-de los Santos, Q.M., Eren, A.M., van der Wielen, P.W.J.J., Ijaz, U.Z., Pinto, A.J., 2020. Differential prevalence and host-association of antimicrobial resistance traits in disinfected and non-disinfected drinking water systems. Sci. Total Environ. 749, 141451. https://doi.org/10.1016/j.scitotenv.2020.141451

Shanthi Amudhan, M., Sekar, U., Kamalanathan, A., Balaraman, S., 2012. blaIMP and blaVIM mediated carbapenem resistance in pseudomonas and acinetobacter species in India. J. Infect. Dev. Ctries. 6, 757–762. https://doi.org/10.3855/jidc.2268

Shi, P., Jia, S., Zhang, X.X., Zhang, T., Cheng, S., Li, A., 2013. Metagenomic insights into chlorination effects on microbial antibiotic resistance in drinking water. Water Res. 47, 111–120. https://doi.org/10.1016/j.watres.2012.09.046

Siguier, P., Perochon, J., Lestrade, L., Mahillon, J., Chandler*, and M., 2006. ISfinder: the reference centre for bacterial insertion sequences. Nucleic Acids Res. 34, D32–D36. https://doi.org/10.1093/nar/gkj014

Smeets, P.W.M.H., Medema, G.J., Van Dijk, J.C., 2009. The Dutch secret: How to provide safe drinking water without chlorine in the Netherlands. Drink. Water Eng. Sci. 2, 1–14. https://doi.org/10.5194/dwes-2-1-2009

Stackelberg, P.E., Furlong, E.T., Meyer, M.T., Zaugg, S.D., Henderson, A.K., Reissman, D.B., 2004. Persistence of pharmaceutical compounds and other organic wastewater contaminants in a conventional drinking-water-treatment plant. Sci. Total Environ. 329, 99–113. https://doi.org/10.1016/j.scitotenv.2004.03.015

Stange, C., Sidhu, J.P.S., Toze, S., Tiehm, A., 2019. Comparative removal of antibiotic resistance genes during chlorination, ozonation, and UV treatment. Int. J. Hyg. Environ. Health 222, 541–548. https://doi.org/10.1016/j.ijheh.2019.02.002

Su, H.C., Liu, Y.S., Pan, C.G., Chen, J., He, L.Y., Ying, G.G., 2018. Persistence of antibiotic resistance genes and bacterial community changes in drinking water treatment system: From drinking water source to tap water. Sci. Total Environ. 616-617, 453–461. https://doi.org/10.1016/j.scitotenv.2017.10.318

Tacconelli, E., Magrini, N., 2017. Global priority list of antibiotic-resistant bacteria to guide research, discovery, and development of new antibiotics., World Health Organization.

Tekerlekopoulou, A.G., Pavlou, S., Vayenas, D. V., 2013. Removal of ammonium, iron and manganese from potable water in biofiltration units: A review. J. Chem. Technol. Biotechnol. 88, 751–773. https://doi.org/10.1002/jctb.4031

Tiwari, A., Hokajärvi, A.M., Domingo, J.S., Elk, M., Jayaprakash, B., Ryu, H., Siponen, S., Vepsäläinen, A., Kauppinen, A., Puurunen, O., Artimo, A., Perkola, N., Huttula, T., Miettinen, I.T., Pitkänen, T., 2021. Bacterial diversity and predicted enzymatic function in a multipurpose surface water system – from wastewater effluent discharges to drinking water production. Environ. Microbiomes 16, 1–17. https://doi.org/10.1186/s40793-021-00379-w

UNEP - United Nations Environment Programme, 2016. A Snapshot of the World ‘ s Water Quality: Towards a global assessment, United Nations Environment Programme.

United Nations, 2018. Water Action Decade [WWW Document]. URL https://wateractiondecade.org/ (accessed 1.26.22).

United Nations, 2015. Sustainable Development Goals [WWW Document]. URL https://www.un.org/sustainabledevelopment/water-and-sanitation/ (accessed 1.26.22).

United States Environmental Protection Agency, 2021. Types of Drinking Water Contaminants [WWW Document]. URL https://www.epa.gov/ccl/types-drinking-water-contaminants (accessed 1.26.22).

van Halem, D., Rietveld, L.C., 2014. Introduction to Drinking Water Treatment [WWW Document]. URL https://online-learning.tudelft.nl/courses/introduction-to-drinking-water-treatment/ (accessed 11.4.20).

Velten, S., Boller, M., Köster, O., Helbing, J., Weilenmann, H.U., Hammes, F., 2011. Development of biomass in a drinking water granular active carbon (GAC) filter. Water Res. 45, 6347–6354. https://doi.org/10.1016/j.watres.2011.09.017

Volkmann, H., Schwartz, T., Bischoff, P., Kirchen, S., Obst, U., 2004. Detection of clinically relevant antibiotic-resistance genes in municipal wastewater using real-time PCR (TaqMan). J. Microbiol. Methods 56, 277–286. https://doi.org/10.1016/j.mimet.2003.10.014

Waak, M.B., Hozalski, R.M., Hallé, C., Lapara, T.M., 2019. Comparison of the microbiomes of two drinking water distribution systems - With and without residual chloramine disinfection. Microbiome 7. https://doi.org/10.1186/s40168-019-0707-5

Wan, K., Guo, L., Ye, C., Zhu, J., Zhang, M., Yu, X., 2021. Accumulation of antibiotic resistance genes in full-scale drinking water biological activated carbon (BAC) filters during backwash cycles. Water Res. 190, 116744. https://doi.org/10.1016/j.watres.2020.116744

Wan, K., Zhang, M., Ye, C., Lin, W., Guo, L., Chen, S., Yu, X., 2019. Organic carbon: An overlooked factor that determines the antibiotic resistome in drinking water sand filter biofilm. Environ. Int. 125, 117–124. https://doi.org/10.1016/j.envint.2019.01.054

Warnecke, F., Amann, R., Pernthaler, J., 2004. Actinobacterial 16S rRNA genes from freshwater habitats cluster in four distinct lineages. Environ. Microbiol. 6, 242–253. https://doi.org/10.1111/J.1462-2920.2004.00561.X

WHO, 2017. Guidelines for the prevention and control of carbapenem-resistant Enterobacteriaceae, Acinetobacter baumannii and Pseudomonas aeruginosa in health care facilities.

Xu, L., Campos, L.C., Canales, M., Ciric, L., 2020. Drinking water biofiltration: Behaviour of antibiotic resistance genes and the association with bacterial community. Water Res. 182, 115954. https://doi.org/10.1016/j.watres.2020.115954

Xu, L., Ouyang, W., Qian, Y., Su, C., Su, J., Chen, H., 2016. High-throughput profiling of antibiotic resistance genes in drinking water treatment plants and distribution systems. Environ. Pollut. 213, 119–126. https://doi.org/10.1016/j.envpol.2016.02.013

Yu, L., Rozemeijer, J., Van Breukelen, B.M., Ouboter, M., Van Der Vlugt, C., Broers, H.P., 2018. Groundwater impacts on surface water quality and nutrient loads in lowland polder catchments: Monitoring the greater Amsterdam area. Hydrol. Earth Syst. Sci. 22, 487–508. https://doi.org/10.5194/hess-22-487-2018

Zhang, H., Chang, F., Shi, P., Ye, L., Zhou, Q., Pan, Y., Li, A., 2019. Antibiotic Resistome Alteration by Different Disinfection Strategies in a Full-Scale Drinking Water Treatment Plant Deciphered by Metagenomic Assembly. Environ. Sci. Technol. 53, 2141–2150. https://doi.org/10.1021/acs.est.8b05907

Zhang, S., Lin, W., Yu, X., 2016. Effects of full-scale advanced water treatment on antibiotic resistance genes in the Yangtze Delta area in China. FEMS Microbiol. Ecol. 92, 1–9. https://doi.org/10.1093/femsec/fiw065

Zhang, T., Lv, K., Lu, Q., Wang, L., Liu, X., 2021. Removal of antibiotic-resistant genes during drinking water treatment: A review. J. Environ. Sci. (China) 104, 415–429. https://doi.org/10.1016/j.jes.2020.12.023

