## Supplementary Material for "Microbiome, resistome and mobilome of chlorine-free drinking water treatment systems"

* Shared first co-authorship

^δ^ Shared co-last authorship

^a^ Delft University of Technology, Delft, The Netherlands

^b^ Dunea, Utility for drinking water and nature conservancy, Plein van de Verenigde Naties 11-15, 2719 EG Zoetermeer, the Netherlands

^c^ Evides Water Company N.V., Schaardijk 150, 3063 NH, Rotterdam, The Netherlands

^d^ Department of Biotechnology and Food Science, Division of Analysis and Control of Microbial Systems, Norwegian University of Science and Technology, Trondheim, Norway

### Supplementary material

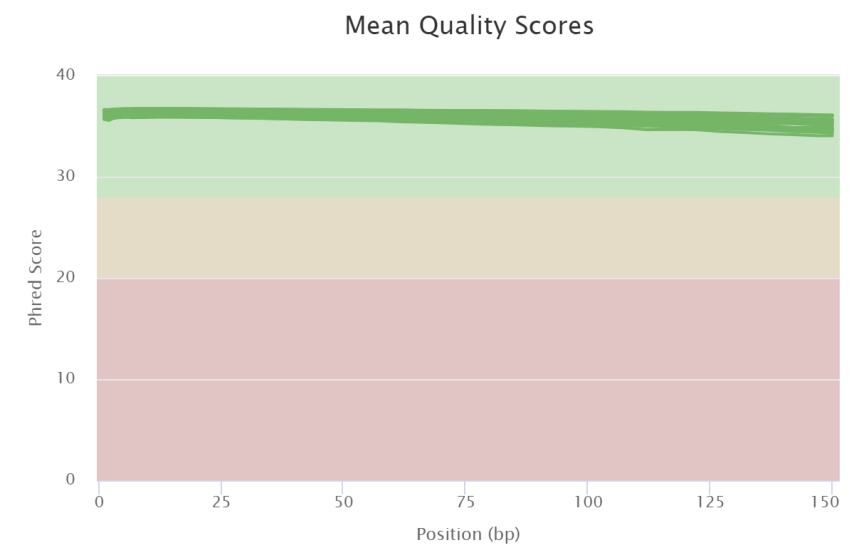

Figure S1. Mean FastQC quality score for the 10 samples sequenced from dune- and reservoir-based DWTPs. Y-axis represents the mean quality score (Phred) obtained by FastQC and X-axis represents the specific position in the read in basepairs (bp).

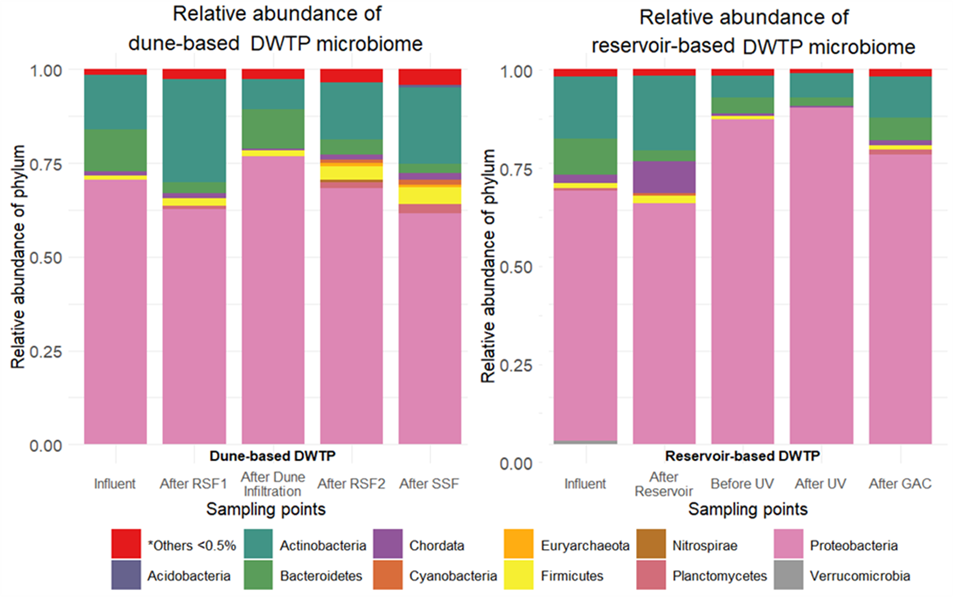

Figure S 2. Microbial community composition at phylum level of dune-based and reservoir-based DWTPs. Relative abundance of the classified phyla (Y-axis) is represented in the different sampling points (X-axis). Phyla with less than 0.5% abundance in all samples was grouped as others.

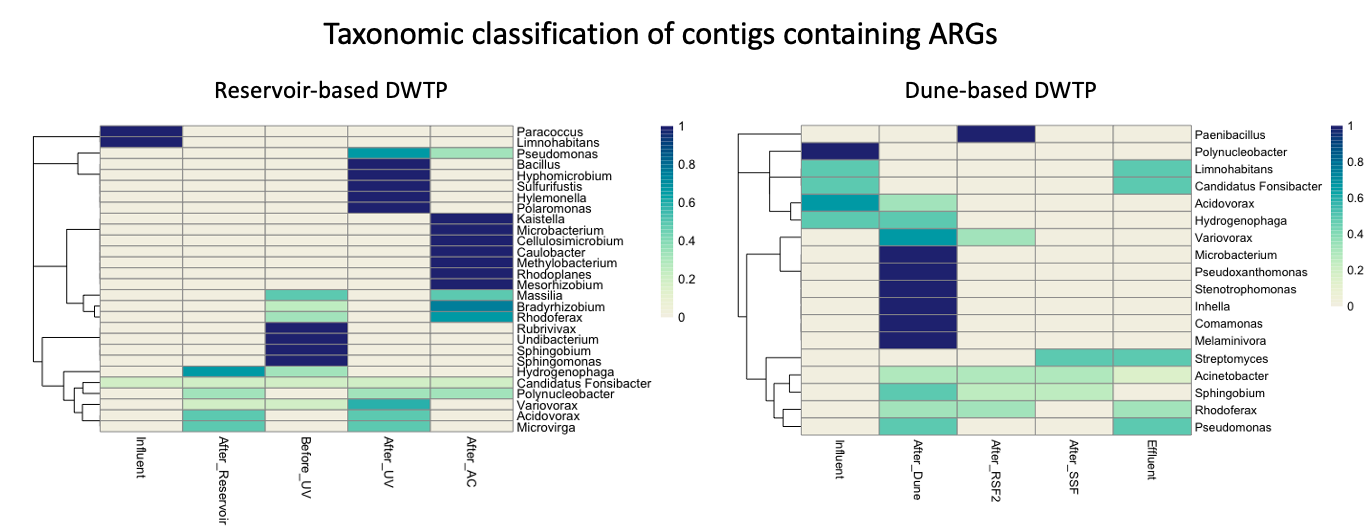

Figure S3. Taxonomic classification at genus-level of contigs encoding ARGs heatmap. Colors represent the relative abundance at genus level in all classified sequences. Labels: (UV) Ultraviolet, (AC) Granular activated carbon, (RSF2) Second Rapid Sand Filtration, (SSF) Slow Sand Filtration.

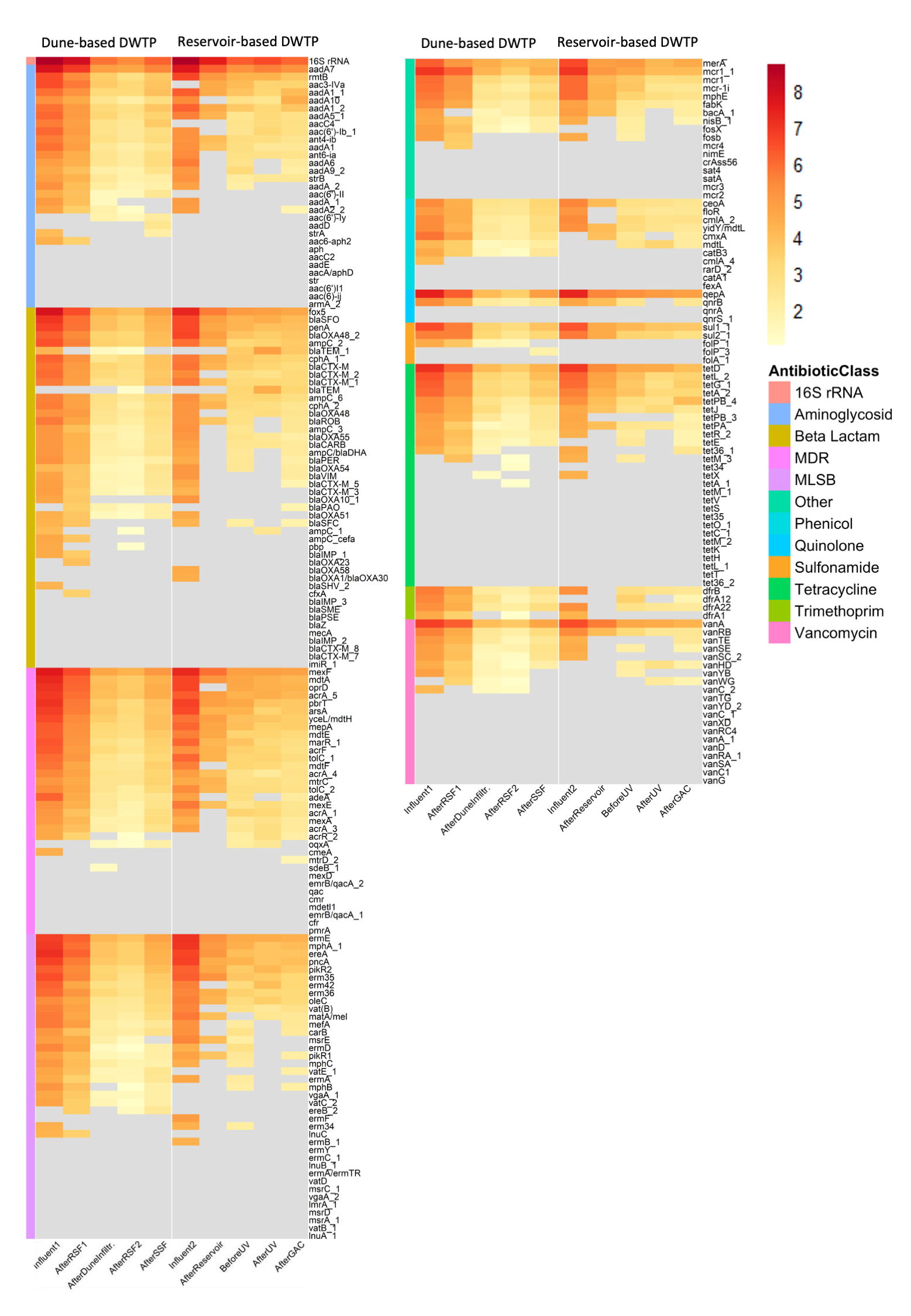

Figure S4. Heatmap of absolute abundance from the 238 ARGs mL^-1^ tested from both dune-based and reservoir-based DWTPs. The absolute abundance of antibiotic is sorted per antibiotic class in logarithmic scale.

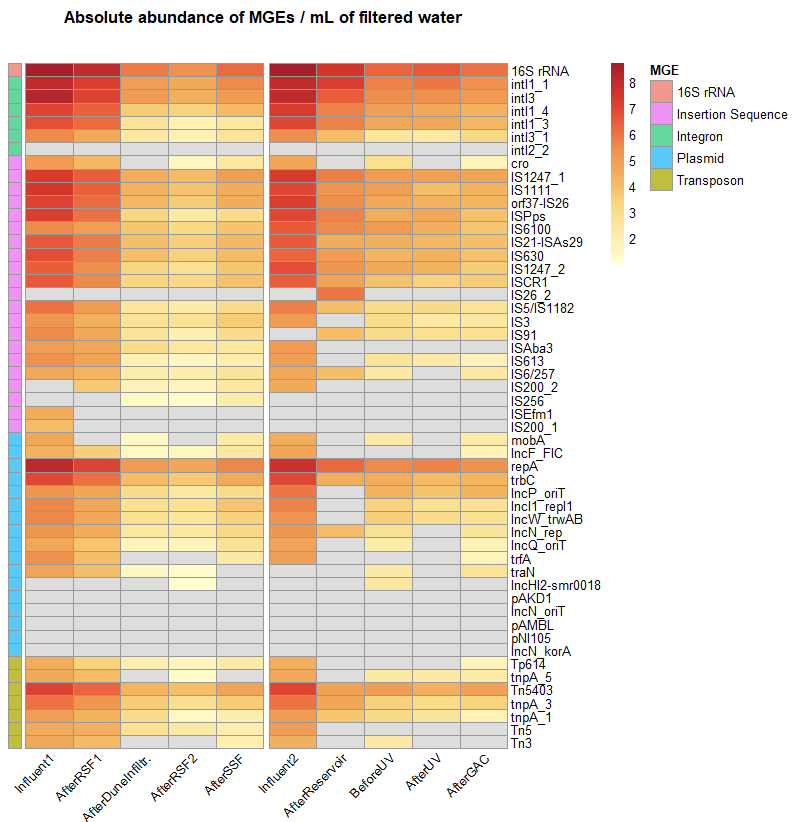

Figure S5. Heatmap of absolute abundance from the MGEs mL^-1^ tested from both dune-based and reservoir-based DWTPs. The absolute abundance of antibiotic is sorted per MGE type in logarithmic scale.

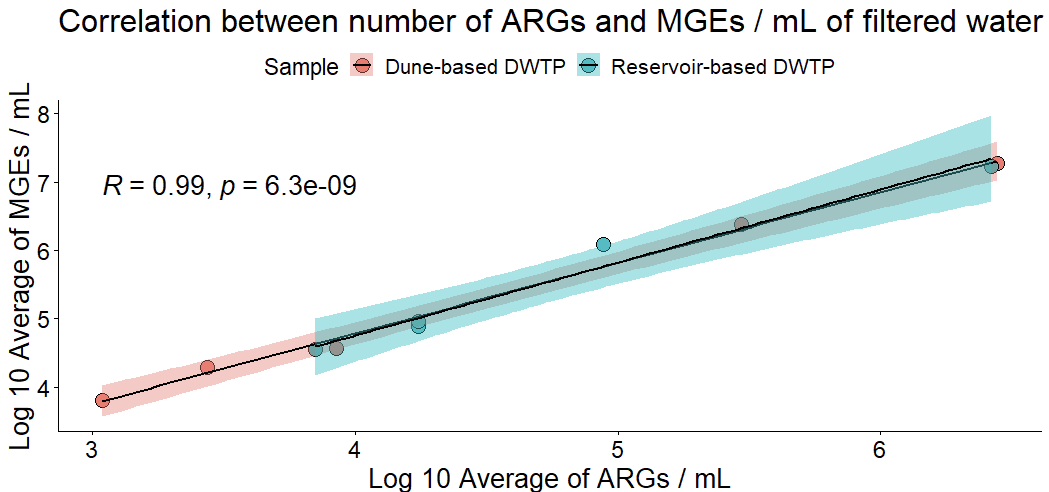

Figure S6. Correlation between absolute abundance of ARGs and MGEs mL^-1^ of filtered water in dune-based and reservoir-based DWTPs. Both axes are displayed in logarithmic scale.

Figure S7. (a) Concentration of ammonia, nitrite and nitrate (mg N mL-1) in dune-based DWTP, including a zoom in ammonia and nitrite concentration. (b) Concentration of ammonia, nitrite and nitrate (mg N mL-1) in reservoir-based DWTP, including a zoom in ammonia and nitrite concentration. (c) Concentration of dissolved organic carbon (DOC) (mg L-1) in dune-based and reservoir-based DWTPs. Values were provided by the DWTPs.

Table S1. Number of raw reads obtained per sample in dune- and reservoir-based DWTPs, specifying the percentages of classified reads, microbial and bacterial percentage (in relation with the percentage of classified reads), number of assembled contigs, and total average length (bp) of assembled contigs per sample. RSF: rapid sand filtration. SSF: slow sand filtration. GAC: granular activated carbon. DWTP: Drinking water treatment plant.

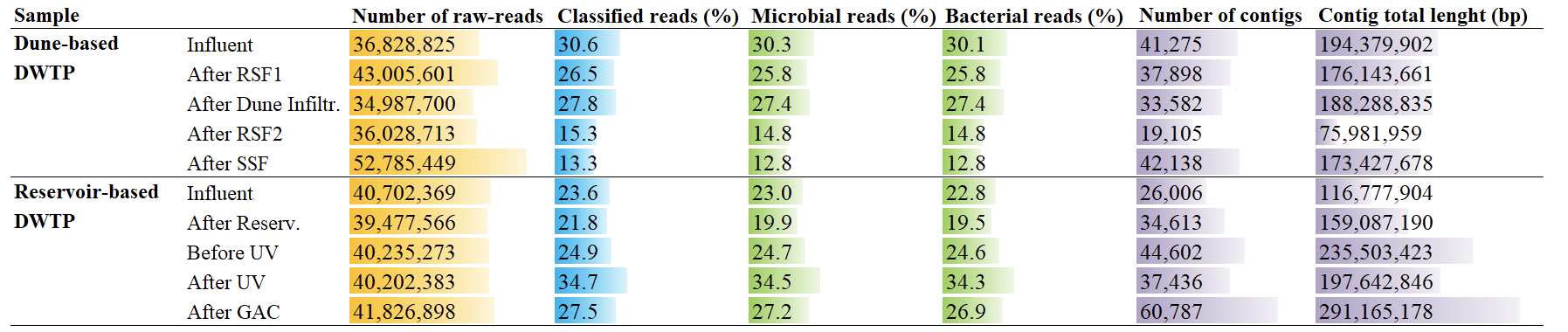

Table S2. List of ARGs analyzed by HT-qPCR. Log removal ARG, MGEs and pathogenic gene copies mL-1 from filtered water coming from dune-based and reservoir-based DWTPs at each sampling point. ARG families include aminoglycoside, beta lactam, MDR, MLSB, other, phenicol, quinolone, sulfonamide, tetracyclin, trimethoprim, and vancomycin. MGEs include integron, IS, transposon and plasmids. Note: MDR: Multi drug resistance; MLSB: macrolide-lincosamide-streptogramin B resistance; IS: Insertion sequence.

|  | | Dune-based DWTP | | | | Reservoir-based DWTP | | | |
| --- | --- | --- | --- | --- | --- | --- | --- | --- | --- |
|  |  | RSF1 | Dune Infiltration | RSF2 | SSF | Reservoir | Before UV | UV disinfection | GAC filtration |
| 16S rRNA ARG | | 0.64 | 2.19 | 0.47 | -0.78 | 1.22 | 1.18 | -0.32 | 0.54 |
| Average of ARGs | | 0.40 | 2.12 | 0.43 | -0.75 | 1.10 | 1.25 | -0.33 | 0.58 |
| Average of MGEs | | 0.88 | 2.05 | 0.47 | -0.75 | 1.04 | 1.22 | -0.07 | 0.43 |
| Average of pathogens | | 0.41 | 0.22 | 0.20 | -0.13 | 4.47 | -1.66 | 0.06 | -0.96 |
| Aminoglyc. | aadA7 | 0.98 | 2.06 | 0.15 | -0.85 | 1.55 | 0.96 | -0.41 | 0.39 |
| Aminoglyc. | rmtB | 1.14 | 1.86 | 0.66 | -0.83 | 1.65 | 0.77 | -0.03 | 0.52 |
| Aminoglyc. | aac3-IVa | 0.79 | 2.17 | 0.40 | -0.63 | -4.50 | 0.44 | 0.75 | -0.07 |
| Aminoglyc. | aadA1_1 | 0.71 | 2.08 | 0.03 | -0.81 | 1.79 | 0.73 | 0.54 | -0.77 |
| Aminoglyc. | aadA10 | 0.75 | 1.96 | 0.53 | -0.82 | 5.46 | -2.76 | 0.25 | -1.95 |
| Aminoglyc. | aadA1_2 | 0.88 | 1.85 | 0.31 | -0.90 | 1.63 | 0.79 | 0.49 | -0.49 |
| Aminoglyc. | aadA5_1 | 1.14 | 1.70 | 0.57 | -0.70 | 1.41 | 1.65 | -0.62 | 0.44 |
| Aminoglyc. | aacC4 | 0.83 | 2.21 | 0.18 | -1.13 | 0.00 | -2.65 | -0.37 | 0.23 |
| Aminoglyc. | aac(6')-Ib_1 | 1.04 | 2.32 | 0.30 | -0.54 | 5.20 | -2.58 | -0.68 | 0.43 |
| Aminoglyc. | ant4-ib | 0.91 | 1.93 | 0.43 | -0.66 | 1.14 | 1.08 | 0.09 | 0.31 |
| Aminoglyc. | aadA1 | 1.00 | 2.49 | 0.44 | -0.60 | 1.46 | 0.89 | 0.56 | -0.03 |
| Aminoglyc. | ant6-ia | 1.00 | 1.84 | 0.37 | -0.76 | 5.36 | -2.83 | 0.07 | 0.12 |
| Aminoglyc. | aadA6 | 0.50 | 1.88 | 0.30 | -0.88 | 5.67 | -2.65 | 2.65 | -2.20 |
| Aminoglyc. | aadA9_2 | 0.67 | 1.97 | 0.14 | -0.84 | 5.08 | -3.13 | 3.13 | -2.02 |
| Aminoglyc. | strB | 0.53 | 1.94 | 0.49 | -0.15 | 5.17 | -2.16 | -0.24 | -0.05 |
| Aminoglyc. | aadA_2 | 1.30 | 1.81 | 0.46 | -0.23 | 4.73 | -1.95 | 1.95 | 0 |
| Aminoglyc. | aac(6')-II | 0.54 | 2.47 | -0.09 | -0.90 | 0 | 0 | 0 | 0 |
| Aminoglyc. | aadA_1 | 1.01 | 2.56 | 1.44 | 0 | 4.95 | 0 | 0 | 0 |
| Aminoglyc. | aadA2_2 | 1.11 | 1.69 | 0.65 | 1.20 | 4.88 | 0 | 0 | -1.73 |
| Aminoglyc. | aac(6')-Iy | 0 | -1.78 | 0.31 | -0.72 | 0 | 0 | 0 | 0 |
| Aminoglyc. | aadD | 0 | 0 | 0 | -2.51 | 0 | 0 | 0 | 0 |
| Aminoglyc. | strA | 4.08 | 0 | 0 | -2.05 | 0 | 0 | 0 | 0 |
| Aminoglyc. | aac6-aph2 | 0.89 | 3.47 | 0 | 0 | 0 | 0 | 0 | 0 |
| Aminoglyc. | aph | 0 | 0 | 0 | 0 | 0 | 0 | 0 | 0 |
| Aminoglyc. | aacC2 | 0 | 0 | 0 | 0 | 0 | 0 | 0 | 0 |
| Aminoglyc. | aadE | 0 | 0 | 0 | 0 | 0 | 0 | 0 | 0 |
| Aminoglyc. | aacA/aphD | 0 | 0 | 0 | 0 | 0 | 0 | 0 | 0 |
| Aminoglyc. | str | 0 | 0 | 0 | 0 | 0 | 0 | 0 | 0 |
| Aminoglyc. | aac(6')I1 | 0 | 0 | 0 | 0 | 0 | 0 | 0 | 0 |
| Aminoglyc. | aac(6)-ij | 0 | 0 | 0 | 0 | 0 | 0 | 0 | 0 |
| Aminoglyc. | armA_2 | 0 | 0 | 0 | 0 | 0 | 0 | 0 | 0 |
| Beta Lactam | fox5 | 1.32 | 2.19 | 0.34 | -0.85 | 1.66 | 0.72 | 0.13 | 0.24 |
| Beta Lactam | blaSFO | 1.04 | 2.01 | 0.36 | -0.80 | 1.76 | 0.66 | -0.27 | 0.40 |
| Beta Lactam | penA | 0.90 | 1.93 | 0.34 | -0.68 | 2.00 | 0.41 | 0.33 | -0.05 |
| Beta Lactam | blaOXA48_2 | 0.74 | 2.10 | 0.16 | -0.87 | 1.80 | 1.01 | 0.33 | -0.23 |
| Beta Lactam | ampC_2 | 0.81 | 2.18 | 0.14 | -0.67 | 1.42 | 1.30 | -0.18 | -0.24 |
| Beta Lactam | blaTEM_1 | 4.23 | -1.37 | 0.25 | 1.13 | 0 | -3.61 | -1.21 | 1.17 |
| Beta Lactam | cphA_1 | 0.94 | 2.69 | -0.33 | -0.95 | 1.32 | 0.73 | 0.26 | 0.14 |
| Beta Lactam | blaCTX-M | 0.90 | 1.71 | 0.26 | -0.86 | 1.61 | 0.62 | 0.28 | 0.32 |
| Beta Lactam | blaCTX-M_2 | 1.10 | 1.75 | 0.28 | -0.84 | 5.40 | -2.91 | 0.28 | -0.47 |
| Beta Lactam | blaCTX-M_1 | 0.94 | 2.05 | 0.04 | -1.05 | 1.88 | 0.73 | -0.07 | 0.14 |
| Beta Lactam | blaTEM | 0 | 0 | -1.12 | 1.12 | 0 | -3.25 | -1.22 | 1.22 |
| Beta Lactam | ampC_6 | 0.81 | 1.91 | 0.45 | -0.81 | 1.37 | 0.69 | -0.12 | 0.45 |
| Beta Lactam | cphA_2 | 0.91 | 2.07 | 0.64 | -0.82 | 0.92 | 1.12 | 0.21 | 0.34 |
| Beta Lactam | blaOXA48 | 0.16 | 2.39 | 0.37 | -0.77 | 5.15 | -2.79 | -0.17 | 0.34 |
| Beta Lactam | blaROB | 1.08 | 2.09 | 0.44 | -0.68 | 1.32 | 0.99 | 2.79 | -2.40 |
| Beta Lactam | ampC_3 | 1.11 | 1.96 | 0.20 | -0.92 | 5.21 | -2.53 | 2.53 | -2.46 |
| Beta Lactam | blaOXA55 | 0.80 | 2.08 | 0.49 | -0.88 | 5.08 | -2.70 | 0.34 | 0.34 |
| Beta Lactam | blaCARB | 0.75 | 2.27 | 0.23 | -0.86 | 4.79 | -2.52 | 0.01 | 0.27 |
| Beta Lactam | ampC/blaDHA | 0.92 | 2.11 | 0.08 | -0.72 | 4.64 | -2.32 | 2.32 | -2.25 |
| Beta Lactam | blaPER | 1.02 | 2.36 | 0.30 | -0.52 | 4.43 | -2.41 | 2.41 | -1.98 |
| Beta Lactam | blaOXA54 | 0.37 | 2.13 | 0.11 | -1.07 | 5.02 | -2.45 | 2.45 | -2.16 |
| Beta Lactam | blaVIM | 0.97 | 2.04 | 0.17 | -0.85 | 5.04 | 0 | 0 | -2.24 |
| Beta Lactam | blaCTX-M_5 | 0.94 | 2.17 | 0.06 | -0.82 | 4.52 | 0 | 0 | -1.74 |
| Beta Lactam | blaCTX-M_3 | 1.35 | 1.85 | 0.36 | -0.94 | 4.21 | 0 | 0 | 0 |
| Beta Lactam | blaOXA10_1 | 1.01 | 3.65 | 0 | 0 | 5.15 | 0 | 0 | 0 |
| Beta Lactam | blaPAO | -3.53 | 1.75 | 0.34 | -0.57 | 0 | 0 | 0 | -1.76 |
| Beta Lactam | blaOXA51 | 0.60 | 2.26 | 0.21 | -0.77 | 4.53 | 0 | 0 | 0 |
| Beta Lactam | blaSFC | 0.44 | 3.85 | 0 | 0 | 4.22 | -2.12 | 2.12 | -1.84 |
| Beta Lactam | ampC_1 | 4.01 | 0 | -1.01 | 1.01 | 0 | 0 | -2.54 | 2.54 |
| Beta Lactam | ampC_cefa | 0.87 | 3.74 | 0 | 0 | 0 | 0 | 0 | 0 |
| Beta Lactam | pbp | 4.55 | 0 | -1.11 | 1.11 | 0 | 0 | 0 | 0 |
| Beta Lactam | blaIMP_1 | 0.93 | 3.61 | 0 | 0 | 0 | 0 | 0 | 0 |
| Beta Lactam | blaOXA23 | -3.89 | 3.89 | 0 | 0 | 0 | 0 | 0 | 0 |
| Beta Lactam | blaOXA58 | 0 | 0 | 0 | 0 | 4.51 | 0 | 0 | 0 |
| Beta Lactam | blaOXA1/blaOXA30 | 0 | 0 | 0 | 0 | 4.42 | 0 | 0 | 0 |
| Beta Lactam | blaSHV_2 | 4.17 | 0 | 0 | 0 | 0 | 0 | 0 | 0 |
| Beta Lactam | cfxA | -3.46 | 3.46 | 0 | 0 | 0 | 0 | 0 | 0 |
| Beta Lactam | blaIMP_3 | 0 | 0 | 0 | 0 | 0 | 0 | 0 | 0 |
| Beta Lactam | blaSME | 0 | 0 | 0 | 0 | 0 | 0 | 0 | 0 |
| Beta Lactam | blaPSE | 0 | 0 | 0 | 0 | 0 | 0 | 0 | 0 |
| Beta Lactam | blaZ | 0 | 0 | 0 | 0 | 0 | 0 | 0 | 0 |
| Beta Lactam | mecA | 0 | 0 | 0 | 0 | 0 | 0 | 0 | 0 |
| Beta Lactam | blaIMP_2 | 0 | 0 | 0 | 0 | 0 | 0 | 0 | 0 |
| Beta Lactam | blaCTX-M_8 | 0 | 0 | 0 | 0 | 0 | 0 | 0 | 0 |
| Beta Lactam | blaCTX-M_7 | 0 | 0 | 0 | 0 | 0 | 0 | 0 | 0 |
| Beta Lactam | imiR_1 | 0 | 0 | 0 | 0 | 0 | 0 | 0 | 0 |
| Integron | intI1_1 | 0.83 | 2.24 | 0.37 | -0.96 | 0.76 | 1.52 | -0.16 | 0.50 |
| Integron | intl3 | 1.13 | 2.03 | 0.66 | -0.59 | 1.67 | 0.93 | 0.13 | 0.19 |
| Integron | intI1_4 | 0.71 | 2.77 | 0.22 | -0.80 | 1.72 | 0.68 | 0.33 | 0.30 |
| Integron | intI1_3 | 0.62 | 3.53 | 0.74 | -0.52 | 1.27 | 1.03 | 0.01 | 0.42 |
| Integron | intI3_1 | 0.91 | 2.24 | 0.48 | -0.76 | 1.38 | 1.60 | 0.10 | -0.85 |
| Integron | intI2_2 | 0 | 0 | 0 | 0 | 0 | 0 | 0 | 0 |
| MDR | mexF | 1.03 | 1.89 | 0.38 | -0.74 | 1.39 | 1.01 | -0.02 | 0.15 |
| MDR | mdtA | 1.20 | 2.24 | 0.28 | -0.71 | 1.58 | 0.46 | 0.28 | 0.27 |
| MDR | oprD | 0.95 | 2.03 | 0.48 | -0.83 | 6.68 | -4.60 | 0.27 | 0.08 |
| MDR | acrA_5 | 0.96 | 1.94 | 0.30 | -0.77 | 1.40 | 0.61 | -0.13 | 0.29 |
| MDR | pbrT | 1.33 | 2.15 | 0.52 | -0.79 | 1.87 | 0.29 | 0.77 | 0.14 |
| MDR | arsA | 1.15 | 1.98 | 0.52 | -0.62 | 2.30 | 0.01 | 0.93 | -0.30 |
| MDR | yceL/mdtH | 1.22 | 1.83 | 0.45 | -0.80 | 1.01 | 0.88 | 0.20 | 0.25 |
| MDR | mepA | 0.95 | 2.09 | 0.27 | -0.99 | 1.37 | 0.75 | 0.50 | -0.04 |
| MDR | mdtE | 0.97 | 2.12 | 0.21 | -0.71 | 0.98 | 1.23 | -0.02 | 0.25 |
| MDR | marR_1 | 1.23 | 2.26 | 0.38 | -0.92 | 1.94 | 0.36 | 0.65 | 0.26 |
| MDR | acrF | 1.30 | 1.88 | 0.43 | -0.69 | 0.97 | 0.83 | 0.17 | 0.45 |
| MDR | tolC_1 | 0.91 | 2.43 | 0.32 | -0.92 | 1.75 | 1.00 | -0.05 | 0.36 |
| MDR | mdtF | 1.17 | 2.09 | 0.12 | -1.01 | 5.58 | -3.28 | 0.11 | 0.51 |
| MDR | acrA_4 | 0.81 | 2.02 | 0.09 | -0.65 | 0.99 | 1.13 | 0.43 | -0.43 |
| MDR | mtrC | 0.64 | 1.93 | -0.02 | -0.76 | 0.33 | 1.38 | -0.05 | 0.19 |
| MDR | tolC_2 | 0.76 | 2.06 | 0.49 | -0.97 | 1.24 | 1.13 | 0.02 | 0.25 |
| MDR | adeA | 1.59 | 2.20 | 0.21 | -0.50 | 4.75 | -2.92 | 2.92 | -2.21 |
| MDR | mexE | 0.92 | 1.95 | 0.60 | -0.86 | 1.87 | 1.30 | -0.43 | 0.51 |
| MDR | acrA_1 | 1.13 | 1.70 | 0.43 | -1.15 | 4.97 | -3.39 | 0.18 | 0.63 |
| MDR | mexA | 0.83 | 2.18 | 0.03 | -1.06 | 4.45 | -2.34 | -0.41 | 0.44 |
| MDR | acrA_3 | 0.67 | 1.91 | 0.23 | -0.89 | 4.79 | -2.48 | -0.44 | 0.64 |
| MDR | acrR_2 | 1.26 | 3.36 | -1.02 | 1.02 | 0 | -2.15 | -0.73 | 0.85 |
| MDR | oqxA | 0 | -1.35 | 0.24 | -0.85 | 0 | -2.16 | -0.23 | 2.39 |
| MDR | cmeA | 4.40 | 0 | 0 | 0 | 0 | 0 | 0 | 0 |
| MDR | mtrD_2 | 0 | 0 | 0 | 0 | 0 | 0 | 0 | -1.67 |
| MDR | sdeB_1 | 0 | -1.26 | 1.26 | 0 | 0 | 0 | 0 | 0 |
| MDR | mexD | 0 | 0 | 0 | 0 | 0 | 0 | 0 | 0 |
| MDR | emrB/qacA_2 | 0 | 0 | 0 | 0 | 0 | 0 | 0 | 0 |
| MDR | qac | 0 | 0 | 0 | 0 | 0 | 0 | 0 | 0 |
| MDR | cmr | 0 | 0 | 0 | 0 | 0 | 0 | 0 | 0 |
| MDR | mdetl1 | 0 | 0 | 0 | 0 | 0 | 0 | 0 | 0 |
| MDR | emrB/qacA_1 | 0 | 0 | 0 | 0 | 0 | 0 | 0 | 0 |
| MDR | cfr | 0 | 0 | 0 | 0 | 0 | 0 | 0 | 0 |
| MDR | pmrA | 0 | 0 | 0 | 0 | 0 | 0 | 0 | 0 |
| IS | cro | 0.83 | 4.22 | -1.47 | -0.95 | 4.77 | -2.80 | 2.80 | -1.65 |
| IS | IS1247_1 | 0.88 | 1.96 | 0.39 | -0.86 | 1.60 | 0.67 | 0.17 | 0.13 |
| IS | IS1111 | 1.06 | 2.03 | 0.50 | -0.68 | 1.71 | 0.31 | 0.98 | -0.37 |
| IS | orf37-IS26 | 1.04 | 1.91 | 0.60 | -0.88 | 1.80 | 0.74 | 0.36 | 0.03 |
| IS | ISPps | 1.12 | 2.79 | 1.00 | -0.70 | 1.32 | 1.18 | -0.16 | 0.78 |
| IS | IS6100 | 0.55 | 1.20 | 0.65 | -0.71 | 1.05 | 0.39 | 0.66 | 0.68 |
| IS | IS21-ISAs29 | 0.82 | 1.79 | 0.50 | -0.72 | 2.00 | 0.06 | 0.26 | 0.20 |
| IS | IS630 | 1.12 | 1.75 | 0.55 | -0.66 | 1.26 | 0.80 | -0.12 | 0.44 |
| IS | IS1247_2 | 0.96 | 2.20 | 0.46 | -0.96 | 1.56 | 0.68 | 0.23 | 0.72 |
| IS | ISCR1 | 1.08 | 2.22 | 0.06 | -0.72 | 1.55 | 0.92 | 0.47 | -0.10 |
| IS | IS26_2 | 0 | 0 | 0 | 0 | -5.88 | 5.88 | 0 | 0 |
| IS | IS5/IS1182 | 1.01 | 2.51 | 0.36 | -0.84 | 1.75 | 0.95 | -0.11 | 0.51 |
| IS | IS3 | 0.83 | 1.77 | -0.06 | -0.90 | 5.07 | -2.94 | 0.66 | -0.06 |
| IS | IS91 | 0.89 | 2.05 | 0.67 | -1.05 | -3.96 | 0.93 | 0.19 | 0.12 |
| IS | ISAba3 | 0.25 | 1.71 | 0.57 | 0.04 | 5.14 | 0 | 0 | 0 |
| IS | IS613 | 0.55 | 3.01 | 0.38 | -0.63 | 5.00 | -2.56 | 0.33 | 0.46 |
| IS | IS6/257 | 0.34 | 2.22 | 0.12 | -0.66 | 0.72 | 1.62 | 2.37 | -2.38 |
| IS | IS200_2 | -3.69 | 1.96 | 0.14 | -0.78 | 4.62 | 0 | 0 | 0 |
| IS | IS256 | 0 | -1.34 | 0.12 | -0.82 | 0 | 0 | 0 | 0 |
| IS | ISEfm1 | 4.49 | 0 | 0 | 0 | 0 | 0 | 0 | 0 |
| IS | IS200_1 | 4.07 | 0 | 0 | 0 | 0 | 0 | 0 | 0 |
| Plasmid | mobA | 4.71 | -1.36 | 1.36 | -2.28 | 4.50 | -2.23 | 2.23 | -2.16 |
| Plasmid | lncF_FIC | 0.87 | 2.13 | -0.19 | -0.89 | 4.72 | 0 | 0 | -1.71 |
| Plasmid | repA | 0.96 | 2.00 | 0.37 | -0.86 | 1.48 | 0.64 | -0.11 | 0.28 |
| Plasmid | trbC | 0.90 | 1.95 | 0.45 | -0.92 | 2.43 | 0.01 | 0.40 | -0.14 |
| Plasmid | IncP_oriT | 0.53 | 1.66 | 0.65 | -0.67 | 5.86 | -4.38 | 0.59 | -0.57 |
| Plasmid | IncI1_repI1 | 0.84 | 1.89 | 0.12 | -1.06 | 5.68 | -3.42 | 0.56 | 0.08 |
| Plasmid | IncW_trwAB | 0.96 | 1.88 | 0.27 | -0.91 | 5.26 | -3.51 | 0.54 | 0.19 |
| Plasmid | IncN_rep | 0.71 | 2.05 | 0.01 | -0.78 | 1.10 | 1.30 | 2.79 | -2.59 |
| Plasmid | IncQ_oriT | 0.89 | 2.14 | 0.07 | -1.10 | 4.80 | -2.13 | 2.13 | -1.78 |
| Plasmid | trfA | 1.37 | 3.97 | 0 | -2.19 | 4.95 | 0 | 0 | -1.70 |
| Plasmid | traN | 0.85 | 2.56 | 0.22 | 1.23 | 0 | -2.24 | 2.24 | -2.69 |
| Plasmid | IncHI2-smr0018 | 0 | 0 | -1.02 | 1.02 | 0 | -2.41 | 2.41 | 0 |
| Plasmid | pAKD1 | 0 | 0 | 0 | 0 | 0 | 0 | 0 | 0 |
| Plasmid | IncN_oriT | 0 | 0 | 0 | 0 | 0 | 0 | 0 | 0 |
| Plasmid | pAMBL | 0 | 0 | 0 | 0 | 0 | 0 | 0 | 0 |
| Plasmid | pNI105 | 0 | 0 | 0 | 0 | 0 | 0 | 0 | 0 |
| Plasmid | IncN_korA | 0 | 0 | 0 | 0 | 0 | 0 | 0 | 0 |
| Transposon | Tp614 | 1.11 | 1.42 | 0.37 | -0.61 | 4.47 | 0 | 0 | -1.66 |
| Transposon | tnpA_5 | 0.51 | 4.07 | -1.23 | 1.23 | 4.64 | -2.27 | 0.02 | 0.08 |
| Transposon | Tn5403 | 0.84 | 2.02 | 0.26 | -0.84 | 2.09 | -0.04 | 0.56 | -0.48 |
| Transposon | tnpA_3 | 0.86 | 1.63 | 0.66 | -0.69 | 1.19 | 1.32 | 0.39 | -0.34 |
| Transposon | tnpA_1 | 0.57 | 1.39 | 1.61 | -0.60 | 1.31 | 0.66 | 0.53 | 0.78 |
| Transposon | Tn5 | 0.15 | 1.67 | 0.58 | 0.24 | 4.95 | 0 | 0 | 0 |
| Transposon | Tn3 | 0.59 | 4.11 | 0 | -1.93 | 4.45 | -2.18 | 2.18 | 0 |
| MLSB | ermE | 0.74 | 2.00 | 0.30 | -0.90 | 1.80 | 0.69 | 0.10 | 0.09 |
| MLSB | mphA_1 | 1.04 | 2.08 | 0.35 | -1.00 | 1.97 | 0.68 | 0.53 | -0.28 |
| MLSB | ereA | 0.99 | 2.34 | 0.37 | -0.84 | 1.54 | 1.20 | 0.14 | -0.01 |
| MLSB | pncA | 1.07 | 2.00 | 0.24 | -0.75 | 2.29 | 0.21 | 0.48 | -0.21 |
| MLSB | pikR2 | 0.78 | 2.15 | 0.28 | -0.96 | 1.50 | 1.01 | -0.50 | 0.47 |
| MLSB | erm35 | 1.00 | 2.13 | 0.65 | -0.84 | 1.01 | 1.46 | 0.10 | 0.46 |
| MLSB | erm42 | 1.51 | 1.80 | 0.22 | -0.95 | 5.61 | -3.08 | -0.90 | 0.78 |
| MLSB | erm36 | 0.84 | 1.93 | 0.55 | -0.76 | 1.17 | 0.97 | 0.22 | 0.05 |
| MLSB | oleC | 0.73 | 2.09 | 0.14 | -0.98 | 1.35 | 1.19 | -0.53 | 0.45 |
| MLSB | vat(B) | 0.79 | 2.39 | 0.10 | -0.92 | 5.50 | -2.83 | 0.44 | -0.31 |
| MLSB | matA/mel | 1.05 | 2.31 | 0.31 | -0.88 | 1.77 | 3.99 | -2.50 | 0.13 |
| MLSB | mefA | 0.88 | 2.74 | 0.30 | -0.51 | 5.20 | -2.17 | 2.17 | -2.38 |
| MLSB | carB | 1.22 | 1.67 | 0.16 | -0.64 | 5.24 | -2.63 | 2.63 | -2.31 |
| MLSB | msrE | 0.38 | 2.56 | 0.66 | 1.33 | 1.46 | 1.96 | 2.06 | 0 |
| MLSB | ermD | 0.86 | 3.24 | 0.35 | -0.95 | 4.33 | -2.78 | 2.78 | 0 |
| MLSB | pikR1 | 0.79 | 2.40 | 0.14 | -0.66 | 0.92 | 1.51 | 2.34 | -2.02 |
| MLSB | mphC | 1.00 | 1.96 | 0.33 | -0.11 | 4.45 | -2.30 | 2.30 | 0 |
| MLSB | vatE_1 | 0.74 | 2.27 | 0 | -0.83 | 0 | 0 | 0 | -2.01 |
| MLSB | ermA | 1.03 | 1.71 | 0.14 | -0.83 | 4.63 | -2.23 | 2.23 | 0 |
| MLSB | mphB | 1.26 | 3.88 | -1.05 | -0.92 | 0 | -2.06 | 2.06 | -2.06 |
| MLSB | vgaA_1 | 0.97 | 2.14 | 0.06 | -0.51 | 0 | 0 | 0 | 0 |
| MLSB | vatC_2 | 1.28 | 1.83 | 0.45 | -0.79 | 0 | 0 | 0 | 0 |
| MLSB | ereB_2 | -3.50 | 3.50 | -1.35 | -1.01 | 0 | 0 | 0 | 0 |
| MLSB | ermF | 0 | 0 | 0 | 0 | 4.99 | 0 | 0 | 0 |
| MLSB | erm34 | 4.38 | 0 | 0 | 0 | 4.38 | -2.15 | 2.15 | 0 |
| MLSB | lnuC | 0.88 | 3.43 | 0 | 0 | 0 | 0 | 0 | 0 |
| MLSB | ermB_1 | 0 | 0 | 0 | 0 | 4.39 | 0 | 0 | 0 |
| MLSB | ermY | 0 | 0 | 0 | 0 | 0 | 0 | 0 | 0 |
| MLSB | ermC_1 | 0 | 0 | 0 | 0 | 0 | 0 | 0 | 0 |
| MLSB | lnuB_1 | 0 | 0 | 0 | 0 | 0 | 0 | 0 | 0 |
| MLSB | ermA/ermTR | 0 | 0 | 0 | 0 | 0 | 0 | 0 | 0 |
| MLSB | vatD | 0 | 0 | 0 | 0 | 0 | 0 | 0 | 0 |
| MLSB | msrC_1 | 0 | 0 | 0 | 0 | 0 | 0 | 0 | 0 |
| MLSB | vgaA_2 | 0 | 0 | 0 | 0 | 0 | 0 | 0 | 0 |
| MLSB | lmrA_1 | 0 | 0 | 0 | 0 | 0 | 0 | 0 | 0 |
| MLSB | msrD | 0 | 0 | 0 | 0 | 0 | 0 | 0 | 0 |
| MLSB | msrA_1 | 0 | 0 | 0 | 0 | 0 | 0 | 0 | 0 |
| MLSB | vatB_1 | 0 | 0 | 0 | 0 | 0 | 0 | 0 | 0 |
| MLSB | lnuA_1 | 0 | 0 | 0 | 0 | 0 | 0 | 0 | 0 |
| Other | merA | 1.15 | 0.60 | 0.44 | -0.11 | 1.73 | -0.11 | 0.84 | -0.39 |
| Other | mcr1_1 | 0.91 | 1.89 | 0.68 | -0.85 | 1.91 | 0.38 | 0.66 | 0.01 |
| Other | mcr1 | 0.99 | 1.99 | 0.53 | -0.88 | 1.83 | 0.44 | 0.79 | -0.14 |
| Other | mcr-1i | 0.81 | 2.55 | 0.42 | -0.78 | 1.74 | 0.55 | 1.24 | 0.05 |
| Other | mphE | 0.81 | 2.32 | 0.14 | -0.83 | 1.73 | 0.92 | 0.49 | 0.20 |
| Other | fabK | 1.03 | 2.25 | 0.28 | -0.76 | 0.81 | 1.42 | 0.22 | 0.10 |
| Other | bacA_1 | 4.77 | -1.64 | -0.35 | -0.77 | 0.87 | 1.63 | 2.38 | -2.16 |
| Other | nisB_1 | 1.04 | 1.73 | 0.38 | -0.60 | 4.46 | -2.09 | 2.09 | -1.75 |
| Other | fosX | 0.89 | 2.29 | 0.39 | -0.80 | 0 | -2.09 | 2.09 | 0 |
| Other | fosb | 1.18 | 3.62 | 0 | 0 | 4.55 | -1.94 | 1.94 | 0 |
| Other | mcr4 | -3.48 | 3.48 | 0 | 0 | 0 | 0 | 0 | 0 |
| Other | nimE | 0 | 0 | 0 | 0 | 0 | 0 | 0 | 0 |
| Other | crAss56 | 0 | 0 | 0 | 0 | 0 | 0 | 0 | 0 |
| Other | sat4 | 0 | 0 | 0 | 0 | 0 | 0 | 0 | 0 |
| Other | satA | 0 | 0 | 0 | 0 | 0 | 0 | 0 | 0 |
| Other | mcr3 | 0 | 0 | 0 | 0 | 0 | 0 | 0 | 0 |
| Other | mcr2 | 0 | 0 | 0 | 0 | 0 | 0 | 0 | 0 |
| Phenicol | ceoA | 1.03 | 1.72 | 0.18 | -0.83 | 1.49 | 1.20 | 0.12 | 0.23 |
| Phenicol | floR | 0.58 | 1.89 | 0.13 | -0.82 | 5.14 | -2.58 | 0.21 | -0.06 |
| Phenicol | cmlA_2 | 0.94 | 2.12 | 0.15 | -0.75 | 5.42 | -3.14 | 0.49 | -0.06 |
| Phenicol | yidY/mdtL | 1.23 | 1.87 | 0.30 | -0.87 | 1.25 | 1.10 | 0.33 | 0.27 |
| Phenicol | cmxA | 1.10 | 2.52 | 0.20 | -0.62 | -3.87 | 1.44 | 2.43 | -2.20 |
| Phenicol | mdtL | 0.59 | 2.15 | -0.17 | -0.55 | 0 | -2.68 | -0.66 | 1.10 |
| Phenicol | catB3 | 1.01 | 1.86 | 0.33 | -1.09 | 0 | 0 | 0 | 0 |
| Phenicol | cmlA_4 | 4.09 | 0 | 0 | 0 | 0 | 0 | 0 | 0 |
| Phenicol | rarD_2 | 0 | 0 | 0 | 0 | 0 | 0 | 0 | 0 |
| Phenicol | catA1 | 0 | 0 | 0 | 0 | 0 | 0 | 0 | 0 |
| Phenicol | fexA | 0 | 0 | 0 | 0 | 0 | 0 | 0 | 0 |
| Quinolone | qepA | 1.20 | 2.12 | 0.59 | -1.42 | 1.86 | 0.23 | 0.34 | 0.16 |
| Quinolone | qnrB | 1.01 | 2.20 | 0.27 | -0.56 | 0.69 | 1.39 | 2.55 | -2.41 |
| Quinolone | qnrA | 0 | 0 | 0 | 0 | 0 | 0 | 0 | 0 |
| Quinolone | qnrS_1 | 0 | 0 | 0 | 0 | 0 | 0 | 0 | 0 |
| Sulfonamide | sul1_1 | 0.84 | 2.29 | 0.43 | -0.86 | 1.80 | 0.43 | 0.38 | 0.05 |
| Sulfonamide | sul2_1 | 0.03 | 2.36 | 0.70 | -0.69 | 1.24 | 1.19 | 0.09 | 0.45 |
| Sulfonamide | folP_1 | 0.73 | 2.17 | 0.10 | 1.48 | 0 | 0 | 0 | 0 |
| Sulfonamide | folP_3 | 0 | 0 | 0 | -1.87 | 0 | 0 | 0 | 0 |
| Sulfonamide | folA_1 | 0 | 0 | 0 | 0 | 0 | 0 | 0 | 0 |
| Taxonomic | A. baumannii | 1.06 | -0.48 | 0.20 | -0.22 | 4.85 | -2.17 | -0.17 | -0.97 |
| Taxonomic | P. aeruginosa | 0.58 | 1.94 | 0.31 | -0.74 | 4.95 | -2.02 | 2.02 | 0 |
| Taxonomic | Enterococci | -4.42 | 4.42 | 0 | 0 | 0 | 0 | 0 | 0 |
| Taxonomic | K. pneumoniae | 0 | 0 | 0 | 0 | 0 | 0 | 0 | 0 |
| Taxonomic | Campylobacter | 0 | 0 | 0 | 0 | 0 | 0 | 0 | 0 |
| Taxonomic | Staphylococci | 0 | 0 | 0 | 0 | 0 | 0 | 0 | 0 |
| Tetracycline | tetD | 1.03 | 2.07 | 0.43 | -0.68 | 1.52 | 0.81 | 0.08 | 0.49 |
| Tetracycline | tetL_2 | 1.01 | 2.16 | 0.26 | -0.95 | 1.28 | 0.88 | 0.88 | -0.35 |
| Tetracycline | tetG_1 | 0.93 | 2.19 | 0.27 | -0.74 | 1.32 | 1.01 | 0.84 | -0.73 |
| Tetracycline | tetA_2 | 0.83 | 2.02 | 0.29 | -0.76 | 1.36 | 0.42 | 0.68 | -0.09 |
| Tetracycline | tetPB_4 | 0.95 | 1.87 | 0.47 | -0.97 | 1.34 | 0.99 | 0.25 | -1.34 |
| Tetracycline | tetJ | 0.96 | 1.47 | 0.96 | -0.56 | 0.70 | 1.07 | -0.34 | 0.72 |
| Tetracycline | tetPB_3 | 0.59 | 1.88 | 0.74 | -0.87 | 4.56 | 0 | 0 | -1.86 |
| Tetracycline | tetPA | 1.19 | 2.12 | -0.17 | -0.79 | 0.61 | 1.41 | 0.10 | 0.50 |
| Tetracycline | tetR_2 | 0.65 | 2.00 | 0.22 | -0.36 | 4.39 | -2.19 | 2.19 | -1.76 |
| Tetracycline | tetE | 0.92 | 2.07 | 0.30 | -0.66 | 0 | -2.38 | 2.38 | -1.72 |
| Tetracycline | tet36_1 | 0.89 | 1.87 | 1.63 | 0 | 4.49 | 0 | 0 | -2.54 |
| Tetracycline | tetM_3 | -3.92 | 3.92 | -1.04 | 1.04 | 4.35 | -2.19 | 2.19 | 0 |
| Tetracycline | tet34 | 0 | 0 | -1.23 | 1.23 | 0 | 0 | 0 | 0 |
| Tetracycline | tetX | 0 | -1.31 | 1.31 | 0 | 4.32 | 0 | 0 | 0 |
| Tetracycline | tetA_1 | 0 | 0 | -1.02 | 1.02 | 0 | 0 | 0 | 0 |
| Tetracycline | tetM_1 | 0 | 0 | 0 | 0 | 0 | 0 | 0 | 0 |
| Tetracycline | tetV | 0 | 0 | 0 | 0 | 0 | 0 | 0 | 0 |
| Tetracycline | tetS | 0 | 0 | 0 | 0 | 0 | 0 | 0 | 0 |
| Tetracycline | tet35 | 0 | 0 | 0 | 0 | 0 | 0 | 0 | 0 |
| Tetracycline | tetO_1 | 0 | 0 | 0 | 0 | 0 | 0 | 0 | 0 |
| Tetracycline | tetC_1 | 0 | 0 | 0 | 0 | 0 | 0 | 0 | 0 |
| Tetracycline | tetM_2 | 0 | 0 | 0 | 0 | 0 | 0 | 0 | 0 |
| Tetracycline | tetK | 0 | 0 | 0 | 0 | 0 | 0 | 0 | 0 |
| Tetracycline | tetH | 0 | 0 | 0 | 0 | 0 | 0 | 0 | 0 |
| Tetracycline | tetL_1 | 0 | 0 | 0 | 0 | 0 | 0 | 0 | 0 |
| Tetracycline | tetT | 0 | 0 | 0 | 0 | 0 | 0 | 0 | 0 |
| Tetracycline | tet36_2 | 0 | 0 | 0 | 0 | 0 | 0 | 0 | 0 |
| Trimethoprim | dfrB | 0.96 | 2.46 | 0.27 | -0.76 | 5.27 | -2.86 | 0.28 | 0.11 |
| Trimethoprim | dfrA12 | 1.22 | 1.59 | 1.03 | -0.60 | 0 | -3.45 | 3.45 | -1.76 |
| Trimethoprim | dfrA22 | 0.87 | 2.17 | 0.16 | -0.81 | 5.28 | -2.64 | 0.20 | 0.15 |
| Trimethoprim | dfrA1 | 1.14 | 3.41 | -1.06 | 1.06 | 4.30 | 0 | 0 | 0 |
| Vancomycin | vanA | 0.60 | 2.19 | 0.34 | -1.00 | 0.88 | 1.07 | 0.09 | 0.22 |
| Vancomycin | vanRB | 1.07 | 2.17 | 0.29 | -0.91 | 1.35 | 0.92 | 0.67 | 0.01 |
| Vancomycin | vanTE | 0.72 | 1.99 | 0.71 | -0.82 | 4.32 | 0 | 0 | -2.36 |
| Vancomycin | vanSE | 0.91 | 2.27 | -0.20 | -0.86 | 4.49 | -1.95 | 1.95 | -1.80 |
| Vancomycin | vanSC_2 | 0.85 | 2.47 | 0.23 | -1.26 | 4.37 | 0 | 0 | 0 |
| Vancomycin | vanHD | 0.94 | 1.79 | 0.49 | -0.80 | 0 | -2.09 | -0.75 | 0.92 |
| Vancomycin | vanYB | 1.19 | 1.99 | 0.14 | 1.48 | 0 | -1.98 | 1.98 | 0 |
| Vancomycin | vanWG | -3.38 | 2.06 | 0.20 | 1.12 | 0 | 0 | -2.29 | 0.57 |
| Vancomycin | vanC_2 | 4.12 | -1.39 | 0.11 | 1.28 | 0 | 0 | 0 | 0 |
| Vancomycin | vanTG | 0 | 0 | 0 | 0 | 0 | 0 | 0 | 0 |
| Vancomycin | vanYD_2 | 0 | 0 | 0 | 0 | 0 | 0 | 0 | 0 |
| Vancomycin | vanC_1 | 0 | 0 | 0 | 0 | 0 | 0 | 0 | 0 |
| Vancomycin | vanXD | 0 | 0 | 0 | 0 | 0 | 0 | 0 | 0 |
| Vancomycin | vanRC4 | 0 | 0 | 0 | 0 | 0 | 0 | 0 | 0 |
| Vancomycin | vanA_1 | 0 | 0 | 0 | 0 | 0 | 0 | 0 | 0 |
| Vancomycin | vanD | 0 | 0 | 0 | 0 | 0 | 0 | 0 | 0 |
| Vancomycin | vanRA_1 | 0 | 0 | 0 | 0 | 0 | 0 | 0 | 0 |
| Vancomycin | vanSA | 0 | 0 | 0 | 0 | 0 | 0 | 0 | 0 |
| Vancomycin | vanC1 | 0 | 0 | 0 | 0 | 0 | 0 | 0 | 0 |
| Vancomycin | vanG | 0 | 0 | 0 | 0 | 0 | 0 | 0 | 0 |

Example of the number of gene copies per volume of filtered water correction. Gene *aadA7* in influent of dune-based DWTP sample

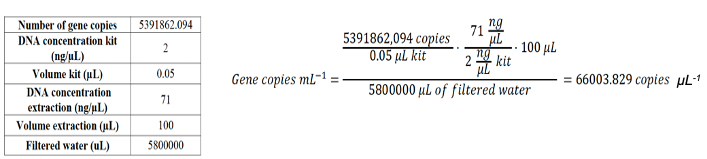

Table S3. Contig nodes with co-occurrence of ARG and MGE. Columns represent the sampling points, the co-occurrence node, the ARGs found, to which antibiotic class they belong, the MGE database that detected the presence of a MGE and the taxonomic classification of such node. The orange color indicates dune-based DWTP results and the blue color indicates reservoir-based DWTP. The MGE database column contains information from plsdb (bacterial plasmids), INTEGRALL (integrons), ISfinder database (bacterial insertion sequences) and ICEberg (bacterial integrative and conjugative elements).

| Sampling point | Node | ARG | Antibiotic class | MGE database | Potential host |
| --- | --- | --- | --- | --- | --- |
| Influent | NODE_11211_length_4463_cov_12.086887 | ant(3'')-Ia_1_X02340 | Aminoglycoside | plsdb, INTEGRALL, ICEberg | Polynucleobacter |
| After RSF1 | NODE_30128_length_2298_cov_17.397236 | ant(3'')-Ia_1_X02340 | Aminoglycoside | plsdb, INTEGRALL, ICEberg | Pseudomonas |
| After RSF1 | NODE_936_length_16827_cov_9.378548 | sul2_2_AY034138 | Sulfonamide | plsdb, INTEGRALL, ICEberg | Acinetobacter |
| After Dune Infiltration | NODE_4_length_434386_cov_14.964421 | blaOXA-287_1_NG_050609 | Beta-lactam | plsdb | Acinetobacter |
| After Dune Infiltration | NODE_1648_length_14981_cov_9.101434 | blaVIM-48_1_KY362199 | Beta-lactam | integrall, IS, ICEberg | Sphingobium |
| After RSF2 | NODE_5007_length_3786_cov_8.195926 | blaVIM-48_1_KY362199 | Beta-lactam | plsdb, INTEGRALL, IS, ICEberg | Sphingobium |
| After SSF | NODE_23294_length_2729_cov_5.609574 | blaVIM-48_1_KY362199 | Beta-lactam | plsdb, INTEGRALL, IS, ICEberg | Sphingobium |
| After Reservoir | NODE_13104_length_3538_cov_23.084410 | ant(3'')-Ia_1_X02340 | Aminoglycoside | plsdb, INTEGRALL, ICEberg | Polynucleobacter |
| Before UV | NODE_324_length_43561_cov_89.313313 | blaVIM-48_1_KY362199 | Beta-lactam | plsdb, INTEGRALL, IS,ICEberg | Sphingobium |
| Before UV | NODE_15748_length_3905_cov_4.388571 | sul1_5_EU780013 | Sulfonamide | plsdb, INTEGRALL, ICEberg | Sphingobium |
| Before UV | NODE_28192_length_2689_cov_3.568717 | blaVIM-48_1_KY362199 | Beta-lactam | plsdb, INTEGRALL, IS, ICEberg | Sphingomonas |
| After UV | NODE_3481_length_9075_cov_11.160089 | blaVIM-48_1_KY362199 | Beta-lactam | plsdb, INTEGRALL, IS, ICEberg | Pseudomonas |
| After UV | NODE_6760_length_6019_cov_9.691147 | blaTEM-181_1_KM977568 | Beta-lactam | plsdb, INTEGRALL, ICEberg | Bacillus |
| After UV | NODE_11087_length_4369_cov_8.523180 | ant(3'')-Ia_1_X02340 | Aminoglycoside | plsdb, INTEGRALL, ICEberg | Polynucleobacter |
| After GAC | NODE_856_length_28072_cov_7.016811 | srm(B)_1_X63451 | Macrolide | plsdb | Rhodoferax |
| After GAC | NODE_15896_length_4245_cov_11.001432 | ant(3'')-Ia_1_X02340 | Aminoglycoside | plsdb, INTEGRALL, ICEberg | Polynucleobacter |
| After GAC | NODE_16870_length_4098_cov_10.446945 | mef(C)_1_AB571865 | Macrolide | plsdb | Kaistella |
| After GAC | NODE_16870_length_4098_cov_10.446945 | mph(G)_1_AB571865 | Macrolide | plsdb | Kaistella |
| After GAC | NODE_49127_length_2249_cov_3.412944 | blaVIM-48_1_KY362199 | Beta-lactam | plsdb, INTEGRALL, IS, ICEberg | Pseudomonas |

Table S 4. List of genes associated with the pathogenic bacteria analyzed by HT-qPCR.

| Microorganism | Gene | Reference |
| --- | --- | --- |
| *Acinetobacter baumanii* | *ompA* | (McConnell et al., 2018) |
| *Pseudomonas aeruginosa* | *gltA* | (Clifford et al., 2012) |
| *Klebsiella pneumoniae* | *ecfX* | (Clifford et al., 2012) |
| *Enterococci* | *23S rRNA* | (He and Jiang, 2009) |
| *Campylobacter* | *16S rRNA* | (Lübeck et al., 2003) |
| *Staphylococci* | *mecA* | (Volkmann et al., 2004) |
